# A boundary-referenced framework for quantifying shoreline-associated movement in white sharks

**DOI:** 10.64898/2026.07.19.739271

**Authors:** Kristian J. Sexton, Elise F. Danko, Nicholas Cheung, Alexandra Eva DiGiacomo, John Sigman, Koppi Kolyvek, Robert W. Holt

**Author notes:** Corresponding Author: **Corresponding Author**, Kristian J. Sexton.

## Abstract

Persistent habitat edges can structure animal movement, yet many analyses treat boundary geometry only implicitly. Advances in aerial platforms, sensor miniaturization, and battery durations now enable high spatial and temporal resolution tracking, creating new opportunities to quantify fine-scale movements. We develop and apply a boundary-referenced framework to quantify movements of white sharks (*Carcharodon carcharias*) off of Cape Cod, Massachusetts, using ∼50 hours of tracking data.

Trajectories were analyzed in a local Cartesian coordinate system and a shoreline-referenced system in which displacement was decomposed into along-shore and cross-shore components. Four complementary metrics quantified path straightness, net and continuous alignment with the shoreline, and unique shoreline traversal. These metrics enabled movement modes to be defined directly from movement relative to the boundary. A parsimonious rules-based classifier recovered visually assigned modes with 88% leave-one-out cross-validated accuracy. Among 90 mode-separated tracks derived from tracks exceeding 15 min, nearly three quarters consisted primarily of sustained along-shore travel, whereas spatially confined movement and nearshore-offshore transitions were less frequent. Boundary-referenced metrics revealed site-level differences in shoreline alignment and space usage across beaches. Findings demonstrate that boundary-referenced metrics recover recurrent, interpretable movement modes in a large predator and provide a general framework for quantifying movements along environmental boundaries.

## Introduction

Understanding how animals move and search within spatially constrained environments is central to movement ecology. As technologies advance, the opportunity to collect positional data at higher temporal and spatial resolutions has increased dramatically [1–6]. These data sets present new opportunities to understand and classify localized behaviors. Many environments including coastal and nearshore systems are structured by persistent physical boundaries such as shorelines, reef edges, or habitat discontinuities that shape movement, orientation, and search strategies, but are often treated implicitly or ignored in formal analyses. The challenges involved are amplified in coastal systems, where movement occurs in the presence of strong spatial structure (e.g., shoreline and sandbar topography) and where behaviors can shift on the order of minutes as animals alternate between directed travel, localized maneuvering, and transitions between nearshore and offshore environments.

Linear habitat features and edges are widely recognized as organizing elements for movement and foraging [7,8]. For predators, travel parallel to edges can facilitate efficient search, maintain proximity to prey-rich zones, and exploit sensory or bathymetric cues [9–11]. Despite this, quantitative frameworks that explicitly treat a boundary as a reference axis, thereby enabling separation of “parallel travel,” “localized maneuvering,” and “transitions in and out of the study area” on a consistent time scale, remain uncommon in empirical studies and this is particularly true for large marine predators. This is in part due to the fact that fine-scale coastal behavior is difficult to observe continuously using conventional telemetry alone, and also because track-level descriptors (e.g., straightness) are often computed over varied durations [12] complicating comparisons among individuals, sites, and studies.

Modern advances in sensor technology and battery endurance provide an opportunity to bridge this gap by enabling direct, high-resolution quantification of predator movement, with tracking durations ranging from minutes to hours [3,12–16]. Drone-based measurements provide the additional benefit of allowing movements to be directly contextualized within habitat structure. However, analysis requires careful handling of scale and continuity as tracks are often intermittent and metrics that do not control for segment duration can conflate true behavioral differences with differences in observation window length [17–19]. A scale-consistent approach that can extract interpretable movement structure from variable-length trajectories is a key requirement.

White sharks (Carcharodon carcharias) provide a compelling system in which to develop and test such an approach with Cape Cod, Massachusetts emerging as an ideal location. Over the past two decades, Cape Cod has become a major seasonal aggregation site for these animals in the western North Atlantic [20,21]. The region’s white sand bottoms and relatively shallow nearshore waters provide an opportunity for continuous aerial observation along an extensive coastline with differing sandbar structure and local context [12,22–24].

We utilize drone-derived white shark trajectories captured along the eastern shore of Cape Cod to understand how nearshore movement is organized relative to the shoreline boundary. We develop a shoreline-referenced framework that decomposes movement into along-shore and cross-shore components and uses scale-consistent path metrics to distinguish recurrent movement modes. We apply this framework to approximately 50 hours of aerial tracking data collected between 2019 and 2022 to evaluate how movement is organized between sustained shoreline-associated travel, localized maneuvering and transitions between nearshore and offshore areas. We also examine whether we can detect differences in localized movement between coastline sections with distinct geomorphic and ecological contexts. Our results demonstrate that white shark movement in this nearshore system is predominantly shoreline-associated, with comparatively limited nearshore–offshore transitions, and that boundary-referenced metrics detect site-specific differences not fully captured by conventional path descriptors.

## Methods

### Data collection and processing

DJI Phantom 4 Pro V2.0 drones with polarizing filters were used for all flights. All flights were conducted in accordance with Federal Aviation Administration regulations and did not require permits [14,25,26]. Data were collected primarily along the three ∼4-km stretches of coastline off the eastern coast of Cape Cod from 2019 to 2022 (Fig. 1) during the summer and fall months coinciding with seasonal residency [14,21,27]. Flights were conducted from either small boats or from town or private beaches with heights above sea level calculated as described in Sexton et al. [14].

**Figure 1.**
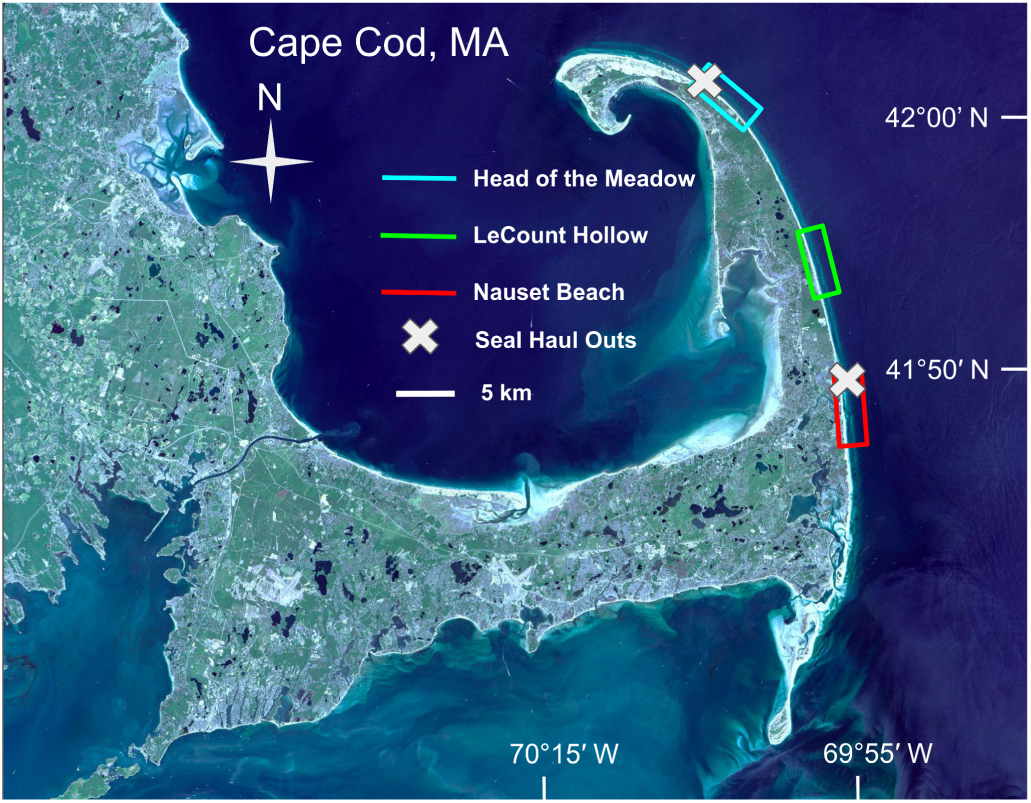
Study area along the outer coast of Cape Cod, Massachusetts, United States. Drone observations of white sharks were conducted at three primary sites: Head of the Meadow in Truro, LeCount Hollow in Wellfleet, and Nauset Beach in Orleans. Colored rectangles indicate the focal survey areas at each beach. Approximate grey seal (Halichoerus grypus) haul-out locations are also shown, as these areas represent key prey resources for white sharks in the region. Satellite image courtesy of NASA Earth Observatory.

At sampling locations, drones were flown along the coast at altitudes between 100 and 122 meters until sharks were spotted. Sharks were subsequently tracked until visual contact was lost, flight limitations were reached, or tracking was discontinued to follow another shark. In instances where visual contact was lost, contributing factors such as glare, turbidity, increased depth, diving, or offshore movement were recorded. The immediate area and that along the animal’s projected path were searched for up to 5 minutes before standard search procedures were re-established. In instances where shark tracking was discontinued due to battery limitations, immediate attempts to re-establish tracking were made following battery changes. In some instances, the use of multiple drones enabled continuous tracking of the same animal across multiple flights.

Data were collected across a range of flight and imaging geometries, including variation in aircraft altitude, gimbal angle, and target position within the field of view (Fig. S1). This variation reflects operational constraints (e.g., glare avoidance) as well as deliberate sampling across varied parameters to support broader downstream uses of data beyond the analyses presented here [14].

All video and flight records were captured, aligned and processed as described in Sexton et al. and detailed in the Supplementary Material, S1.1 [14]. Video data were annotated using a combination of manual and automated processes [14]. Target geographic coordinates were calculated using trigonometric relationships and flight data, including target position on the image sensor, target height relative to the takeoff datum, and known imaging system characteristics [14]. Flight and video data alignment, altitude corrections and heading corrections were implemented using the methods described in Sexton et al. [14].

Shark lengths were measured following the procedures described in Sexton et al. [14]. Speeds were calculated from filtered position and time data with raw speed estimates further filtered using a rolling median absolute deviation filter. Tailbeat frequencies were manually recorded during annotation. Additional details on length, speed, and tailbeat measurements are provided in the Supplementary Material, S1.2. Speed and tailbeat frequency were averaged across entire tracks and reported only for tracks containing >1 min of speed data and >0.5 min of tailbeat-frequency data. Shark positions and directions relative to the shoreline were determined by regularly mapping the coastline with the drone flown directly over the shoreline (Supplementary Material, S1.3, Fig. S2).

### Track grouping, filtering and shoreline transformation

Following the calculation of geographical coordinates for all shark positions, data were grouped into individual tracks, with each track representing the movement of what was believed to be a single shark. Temporary losses of visual contact within a tracking encounter were evaluated as described in the Supplementary Material (S1.4) to determine whether subsequent observations represented a continuation of the same track. Individuals could not generally be reidentified across tracking encounters conducted on different dates or at different locations; tracks were therefore treated as observational units, and the same shark may have contributed more than one track.

Shark and shoreline positions were transformed from geographic coordinates (WGS84) to a local planar coordinate system in meters using a Universal Transverse Mercator (UTM) projection appropriate for the study region. Coordinates were expressed relative to a shared local origin so that shark tracks and shoreline geometry were represented in a common Cartesian (X,Y) reference frame. Shark tracks were filtered as described in Sexton et al. [14], with position accuracy estimated at 11.4 m (95% CI).

For each filtered shark position, the nearest point on the corresponding shoreline polyline was first calculated as a target (Supplementary Material, S1.3). To reduce abrupt jumps in nearest-point associations, movement of the accepted shoreline point was step-limited within continuous shark-track sections. The first valid position in each section was assigned directly to its nearest shoreline point; for each subsequent position, the accepted shoreline point was allowed to move along the shoreline toward the nearest-point target by no more than twice the shark’s displacement between successive positions. The constraint was reset at track discontinuities. The local shoreline bearing at each accepted shoreline location (*shore_dir_i_*) was used as the reference direction (Supplementary Material, S1.3.1).

For successive shark positions (*X_i_*, *Y_i_*), step displacements were calculated as Δ*X_i_*=*X_i_*−*X*_(*i*-1)_ and Δ*Y_i_*=*Y_i_*−*Y*_(*i*-1)_, and step direction was computed as ɸ*_i_*=*atan*2(Δ*Y_i_*, Δ*X_i_*). Step displacements were then rotated into a boundary-aligned frame by subtracting the local shoreline bearing (Fig. 2): 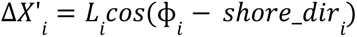 and 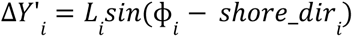, where *L_i_* is step length.

**Figure 2.**
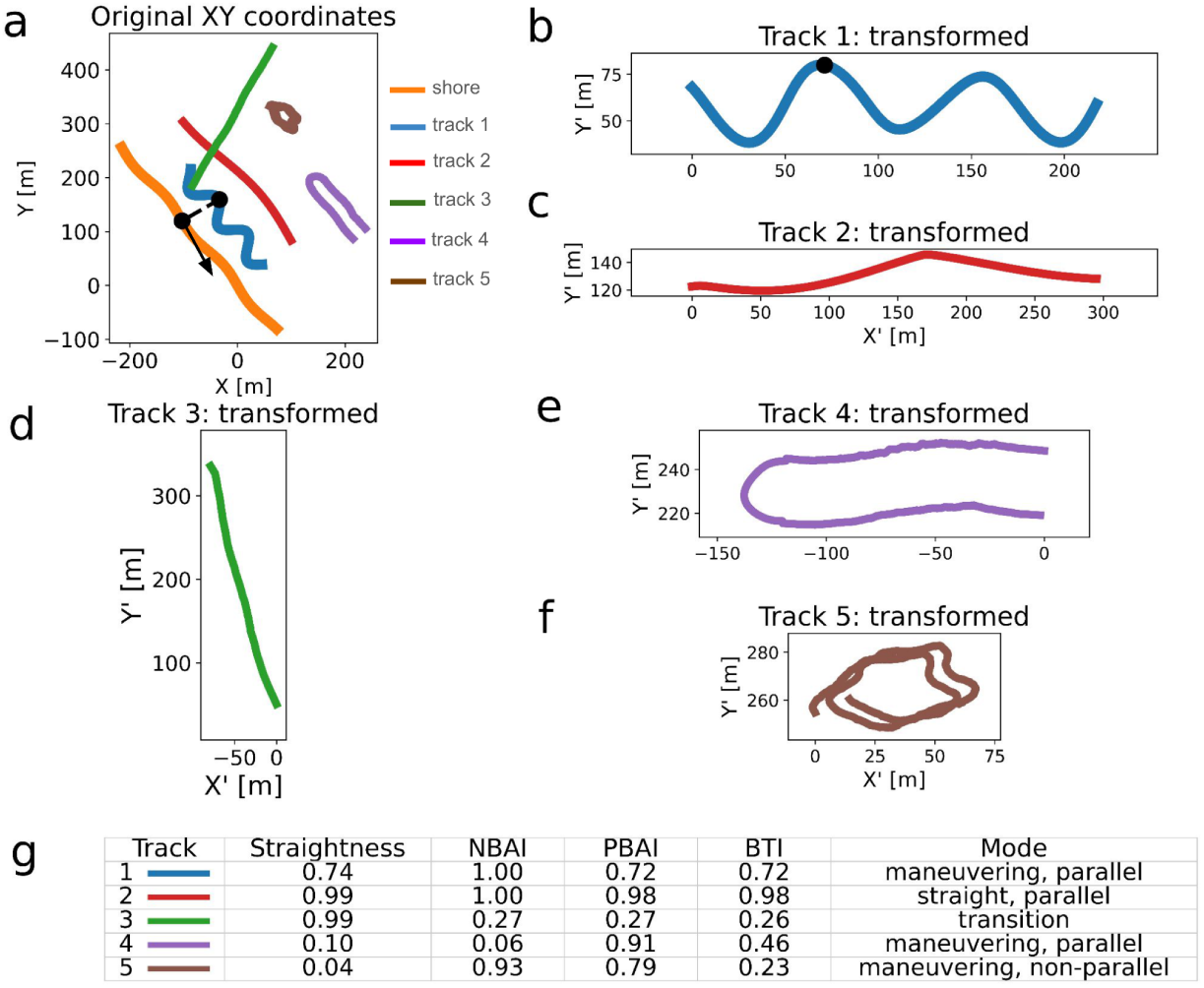
Transformation of standardized example trajectories into a boundary-aligned coordinate system. Each trajectory represents a single 5-min segment with movement speed fixed at 1.0 m s⁻¹. Straightness quantifies path tortuosity in the original reference frame. NBAI captures the overall directional progression of trajectories relative to the boundary. PBAI reflects the instantaneous alignment of movement steps and therefore may decrease when animals follow oscillatory or maneuvering paths that nevertheless progress predominantly along the boundary. BTI quantifies the extent of unique shoreline traversed. (a) Shark trajectories in the original Cartesian coordinate system (X, Y) with the mapped shoreline. A representative shark position on the sinusoidal shark track (blue track) is highlighted (black point), along with its associated shoreline location (black point). The dashed line indicates the offset between the shark position and its associated shoreline location, while the arrow indicates the local shoreline direction at that point. (b–f) The same shark trajectories shown in the boundary-aligned coordinate reference frame. The highlighted point in panel b corresponds to the same shark position shown in panel a. Movement is represented relative to shoreline orientation, with X′ and Y′ representing cumulative along-shore and cross-shore displacement, respectively, and Y′ initialized at the shark’s starting distance from shore. For the approximately linear shoreline analyzed here, this provides an effective boundary-aligned representation of movement. The shoreline is shown at Y′ = 0 as the reference boundary, allowing along-shore and cross-shore movement to be visualized directly. (g) Calculated metrics for each of the five example tracks and the resulting movement modes determined by the rules-based classifier.

The rotated step displacements Δ*X′_i_* and Δ*Y′_i_* represent the along-shore and cross-shore components of movement step *i*, respectively. Cumulative transformed coordinates were calculated as 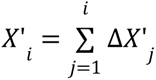 and 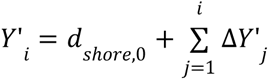 with the start of each track defined as *X*’_0_=0 and *Y*’_0_=*d_s_*_ℎ*ore*,0_ where *d_s_*_ℎ*ore*,0_ is the shark’s initial distance from shore. Thus, Δ*X′_i_* and Δ*Y′_i_* describe individual movement steps, whereas *X′_i_* and *Y′_i_* describe cumulative position in the boundary-aligned reference frame. All shoreline-referenced movement metrics were computed from these rotated step displacements and cumulative transformed coordinates. The shoreline-aligned frame also enabled direct calculation of heading relative to the shoreline.

### Track and segment analysis

Tracks were partitioned into contiguous segments by splitting at temporal gaps >30 s and excluding segments shorter than 5 min. This ensured consistency and enabled the deliberate choice of a relevant time scale despite variations in track length and data continuity. In addition to straightness, we defined three boundary-referenced movement metrics to quantify alignment with, and traversal along, the shoreline: net boundary alignment index (NBAI), path boundary alignment index (PBAI), and boundary traversal index (BTI). All metrics were calculated using overlapping 5-min segments advanced in 15-s increments.

To obtain values across all positions in each track, segment-level metrics were assigned to positions within each segment and averaged across overlapping segments, yielding a continuous representation of movement properties analogous to a centered moving-window approach (Supplementary Material, S1.5) [18,28].

Straightness was calculated as the ratio of net displacement to total path length [29,30] and was computed in the original planar coordinate system where it describes path tortuosity in geographic space [17,31].

NBAI provides a measure of the extent to which a shark’s overall displacement across a track segment is aligned with the shoreline (Fig. 2). The index was calculated as:

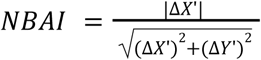

where Δ*X*’ and Δ*Y*’ represent the net displacements along and perpendicular to the shoreline, respectively.

PBAI provides a measure of the extent to which a shark’s movement remains aligned with the shoreline over the full segment (Fig. 2). The index was calculated as:

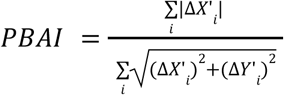

where Δ*X′_i_* and Δ*Y′_i_* are the stepwise displacements along and perpendicular to the shoreline.

BTI quantifies the extent of unique shoreline traversed within each five-minute window (Fig. 2). It was calculated from:

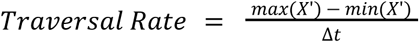

where *X′_i_* is cumulative along-shore position and Δ*t* is the window duration in seconds. This rate *m s*^-1^) was then normalized by the shark’s mean ground speed over the full track to yield a dimensionless BTI.

### Distance from shore and direction of travel

Distance from shore was calculated as the distance between each filtered shark position and its corresponding shoreline position. Distance-from-shore distributions were summarized by converting each valid 10 Hz distance sample to 0.1 s of observation time and aggregating observation time across distance bins. Linear fits to binned observation time were used descriptively to summarize nearshore distance patterns.

Headings were calculated from the direction of movement between successive filtered shark positions. True headings were calculated in the original planar coordinate system, whereas headings relative to the shoreline were calculated in the boundary-aligned reference frame. Both were reported for all available data. A ±30° window was used to define approximately shoreline-parallel movement. Primary direction of travel was calculated using 5-min segments and was defined as the largest 10° bin from the distribution of instantaneous headings within each segment.

### Visual classification of movement modes

Categorical movement-mode assignment requires sufficient observation time and track continuity. All shark tracks over 5 minutes in total duration were categorized into at least one of four movement modes when track continuity and visual interpretability allowed: straight, parallel; maneuvering, parallel; maneuvering, non-parallel and transition (movement between nearshore and offshore environments).

Tracks with more than one clearly distinct mode were considered multi-modal and separated into two or more single-mode tracks. Unless otherwise specified, ‘tracks’ hereafter refers to these final mode-separated tracks. Separation and categorization of tracks were done through visual assessment of tracks alongside the shoreline with all tracks plotted at the same scale. As categorical movement-mode assignment may be sensitive to observation duration, we examined mode composition across several minimum-duration thresholds based on the total duration of initial tracks.

Considering that minimum-duration thresholds may exclude short tracks that end as the result of nearshore–offshore transitions, we categorized all tracks as either potential transitions or non-transitions based on track geometry and the recorded reason that visual contact was lost. Any tracks where at least ¾ of movement in the initial 30 s of the track was directed onshore within 30° of perpendicular to the shoreline were classified as a potential transition. Any track where environmental conditions ultimately resulted in loss of visual contact with the shark and at least ¾ of the final 30 s of the track was directed offshore within 30° of perpendicular was also classified as a potential transition. Additionally, any track where the shark was lost as the result of the animal diving was classified as a potential transition. As a descriptive estimate of continuous inshore activity, total tracking time was divided by the number of potential transitions.

Circling and slinky swim were identified as distinctive movement-pattern subsets during visual assessment. Circling was defined by repeated localized circular movement and was treated as a subset of maneuvering, non-parallel movement. Slinky swim was defined by repeated looping or oscillatory movement embedded within an overall shoreline-aligned trajectory and was treated as a subset of maneuvering, parallel movement. Turn radii during circling and slinky swim were calculated from interactive visualizations of each animal’s track in the original reference frame. Pairs of points along the track were selected to determine distances, allowing the range of turn radii to be derived [14].

Ground speed and tailbeat frequency were compared across visually classified movement modes and subsets, including circling and slinky swim for which these subsets were excluded from their parent modes. Pairwise comparisons were evaluated using Welch’s t-tests with Bonferroni corrections applied separately within each response variable.

### Classification of movement modes

A rules-based classification system was developed based on visual assessment of the relationship between calculated metrics (straightness, NBAI, PBAI, and BTI) and associated modes. All tracks that met continuity requirements for segment analysis and had modes determined based on visual assessment were included. In the simplest version, multi-modal tracks were separated based on visual assessment prior to the determination of track metrics and metrics were averaged across the duration of each track. A decision tree utilizing six conditional if-then statements was used to separate all tracks into one of four distinct movement modes based on straightness, NBAI, PBAI and BTI (Fig. S3).

Decision tree thresholds were optimized algorithmically. Classification performance was evaluated using leave-one-out cross-validation (LOOCV) with fold-specific threshold refitting and bootstrap resampling with out-of-bag testing (Supplementary Methods S1.6).

A dynamic version of the rules-based system applied classification to all eligible points within segments, with final track-level mode based on the most frequent assignment. Multi-modal tracks were split only after metrics were computed.

For comparison purposes a supervised machine-learning classifier (random forest) was also implemented for the determination of movement mode. This classifier utilized the same input metrics (straightness, NBAI, PBAI, and BTI) as the rules-based classifier. Details of model training and validation are provided in the Supplementary Material (Section S1.7).

To enable direct comparison across classification approaches, the dynamic rules-based and random forest classifiers were evaluated on the same 131 mode-separated tracks used by the track-averaged rules-based classifier, each of which independently met segment-analysis requirements after visual separation.

### Comparisons across beach locations

To evaluate whether shoreline-associated movement differed among coastline sections, movement metrics, heading directionality, and mode distributions were compared among beaches. Beach-level differences in boundary-referenced metrics were evaluated using all available track-level summaries meeting segment-analysis requirements, with visually separated modes from multimodal tracks treated as separate units. For each track, directional alignment was summarized as the fraction of valid shoreline-relative headings within ±30° of either shoreline-parallel direction. Straightness, NBAI, PBAI, BTI and track-level shoreline-parallel fractions were compared among beaches using Kruskal–Wallis tests, with Bonferroni-corrected pairwise comparisons and Cliff’s delta effect sizes. Parallel versus maneuvering, non-parallel mode counts were compared among beaches for the mode-separated visually classified tracks derived from initial tracks exceeding 15 min in duration using chi-square contingency-table tests. Pooled directional histograms were treated as descriptive time-use distributions as sequential headings within tracks are autocorrelated.

## Results

### Shark lengths, ground speeds, tailbeat frequencies

Length measurements were obtained from 241 separate shark tracks with error estimates of ±4% (SD) [14]. Total lengths (TL) ranged from 1.97 to 4.63 m (mean ± SD: 3.30 ± 0.44 m; median: 3.35 m) (Fig. S4a) (Supplementary Material, S3.1, S3.2, Fig. S5). Average ground speeds, calculated for 288 tracks, ranged from 0.29–1.81 m s⁻¹ (mean: 1.04 ± 0.27 m s⁻¹; median: 1.04 m s⁻¹) (Fig. S4b). Average tailbeat frequency, measured for 88 tracks, ranged from 0.24–0.63 Hz (mean: 0.43 ± 0.09 Hz; median: 0.43 Hz) (Fig. S4c). All three metrics were collected for 79 tracks. Ground speed scaled linearly with the product of length and tailbeat frequency (Fig. S4d).

### Tracking summary and movement metrics

A total of 359 initial tracks were recorded with durations from seconds to several hours and totaling 50.4 hours of active tracking data and ∼68 hours of total tracking time (Fig. S6a). Total tracking represents the time from the start of tracking to the end of tracking and may include gaps in which shark position was unknown whereas active tracking represents the total time for which shark position could be determined from the aircraft. Data from 121 initial tracks (131 tracks after mode separation) totaling 34.9 hours of active tracking met time and continuity requirements for segment analysis (Fig. S6b). Only these mode-separated tracks were used for metric-based analysis and movement classification (Table S1).

Metrics were calculated dynamically across segments (Fig. 3), with average values for straightness, NBAI, PBAI, and BTI of 0.77, 0.92, 0.84, and 0.75 respectively (Fig. S7a-d). As NBAI is based on net displacement, it captures the overall directional progression of trajectories relative to the shoreline. In contrast, PBAI reflects the instantaneous alignment of movement steps and therefore decreases when animals follow oscillatory or maneuvering paths that nevertheless progress predominantly alongshore (Fig. 2 and Fig. 3). Average straightness, NBAI, PBAI, and BTI varied with visually assessed mode (Fig. S7 and S8) and plotting across tracks revealed clustering, supporting the development of a rule-based classifier (Fig. S3).

**Figure 3.**
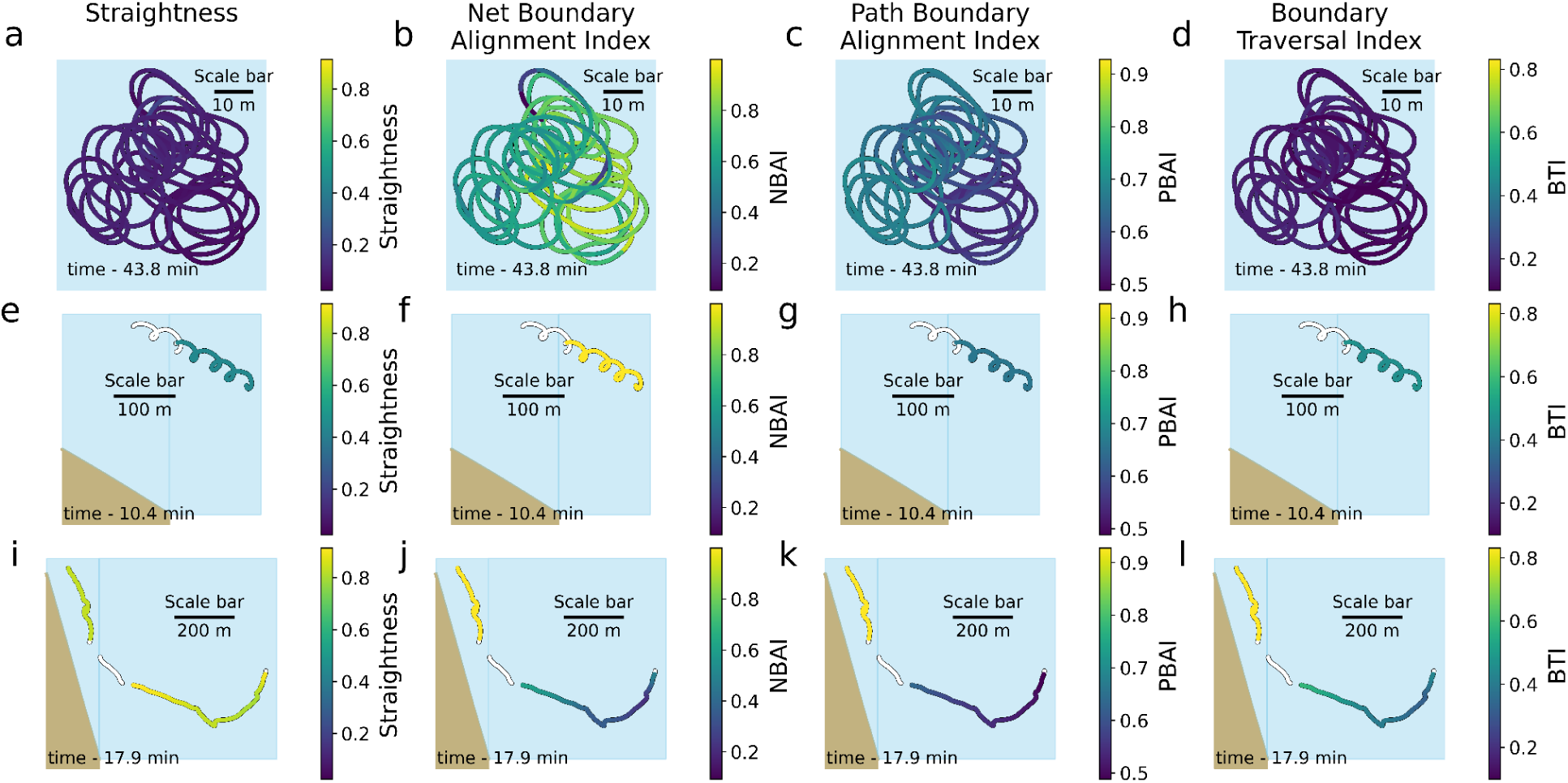
Example tracks illustrating movement metrics used for classification with track color indicating metric value at each position. Straightness, net boundary alignment index, path boundary alignment index and boundary traversal index are plotted for three representative tracks. (a–d) Circling behavior, a subset of maneuvering, non-parallel. (e–h) Slinky swim, a subset of maneuvering, parallel. (i–l) A multimodal track containing both maneuvering, parallel and transition mode. Portions of tracks showing an outline only did not meet the continuity requirements for segment analysis and so metrics are not shown at those positions.

### Movement modes and nearshore durations

Visual classification of the 359 initial tracks produced 229 mode-separated tracks derived from initial tracks exceeding 5 min in total duration (Fig. 4, Table S2). Movement-mode distributions varied with minimum initial-track duration, with the relative frequency of maneuvering, non-parallel tracks increasing at longer duration thresholds (Fig. 4g). With a 15-min threshold (n=90), 37% of tracks were straight, parallel; 36% maneuvering, parallel; 22% maneuvering, non-parallel; and 6% in transition with parallel modes together accounting for 72% of tracks. At 10 minutes (n=149), these were 43%, 34%, 16%, and 7%, respectively with parallel modes together accounting for 77% of tracks.

**Figure 4.**
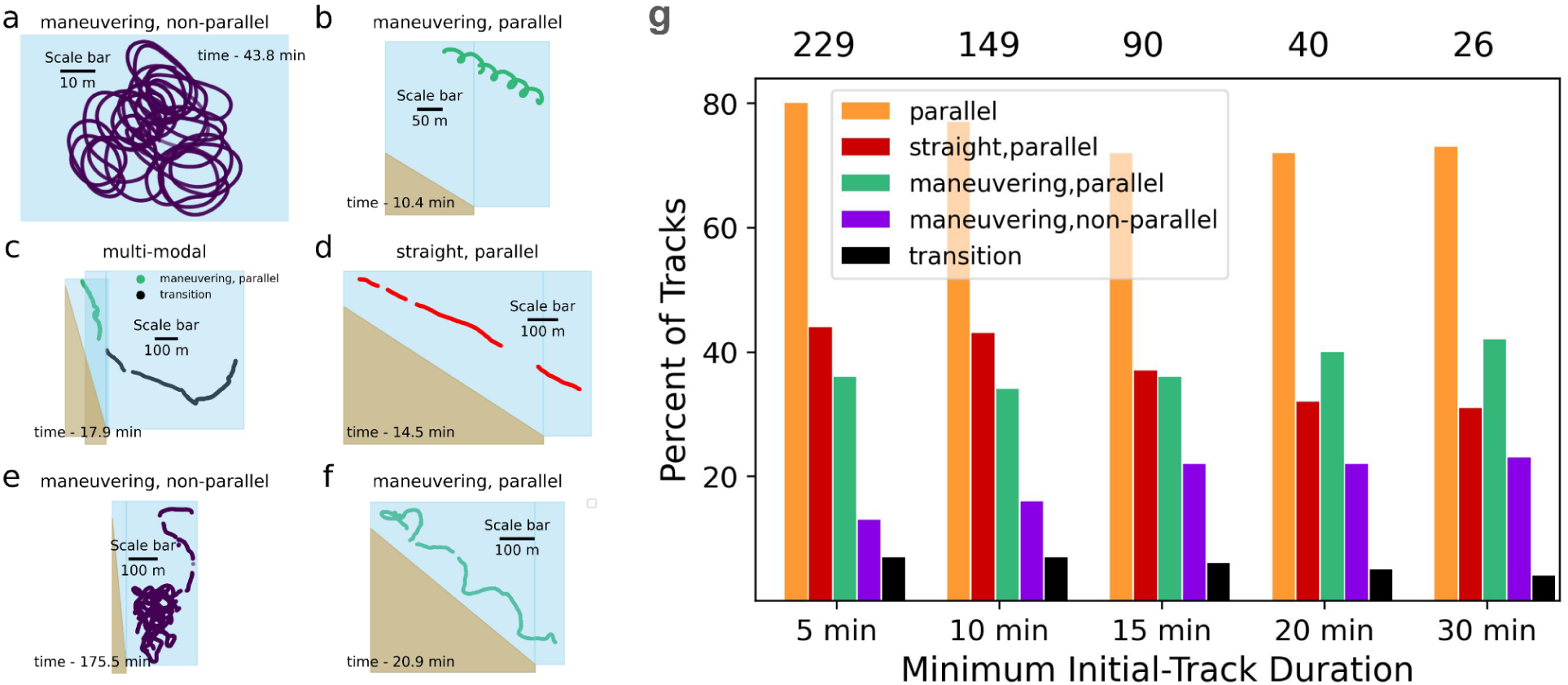
Example animal tracks displaying the movement modes observed, along with the distribution of those modes across a range of minimum track durations. (a) Circling behavior is observed approximately 280 m from shore and represents a subset of the maneuvering, non-parallel mode. (b) Slinky swim, consisting of looping movement embedded within shoreline-parallel travel, is categorized as maneuvering, parallel. (c) The initial portion of the track is categorized as maneuvering, parallel and the shark is then seen to switch to the transition mode. (d) A clear instance of a shark in the straight, parallel mode. (e) The animal maneuvers within a limited area and is categorized as maneuvering, non-parallel. (f) The track displays an overall trajectory parallel to the shoreline but shows oscillation around that trajectory and is categorized as maneuvering, parallel. (g) Comparison of minimum initial track duration with visual classification of movement modes. Numbers above bars indicate the number of mode-separated tracks retained at each initial-track duration threshold. The percentage of each mode, as well as the combined percentage of the two parallel modes, is shown across varying minimum duration thresholds. The proportion of straight, parallel tracks declines with increasing duration, while the total percentage of parallel tracks stabilizes beyond 15-min. The proportion of maneuvering, non-parallel tracks increases up to the 15-min threshold, suggesting that shorter tracks may under-represent this mode due to limited observation time. Minimum track durations in the vicinity of 15-min may improve discrimination between parallel and non-parallel movement.

Potential transitions were identified in 48 of 359 tracks (13%). With ∼68 h of total tracking, this yielded an average of ∼1.4 h of inshore activity per potential transition (Supplementary Material, S3.4). This estimate was consistent with the observation of multiple long-duration tracks lasting on the order of hours (Supplementary Material, Note S4, S4.1, S4.2).

### Movement mode classification performance

Of the 229 visually classified mode-separated tracks derived from initial tracks exceeding 5 min, 131 met continuity requirements for segment analysis and were classified using the rules-based system. Thresholds were selected to formalize visually distinct geometric clusters and were optimized algorithmically using the full set of visually classified tracks. The final threshold set showed 90% agreement with visual classifications (118/131; Fig. S9). To assess generalization performance, leave-one-out cross-validation (LOOCV) was then performed with thresholds refit at each iteration. LOOCV accuracy was 0.88 (115/131), with a balanced accuracy of 0.90.

Bootstrap resampling with out-of-bag testing (1,000 replicates) produced a mean accuracy of 0.84, with a 95% bootstrap interval of 0.73–0.93, and a mean balanced accuracy of 0.87, indicating generally consistent predictive performance across resampled datasets. Misclassifications occurred primarily between straight, parallel and maneuvering, parallel modes. Dynamic classification at the point level yielded slightly lower agreement: 0.76 accuracy at the point level and 0.86 per track, with most errors between straight, parallel and maneuvering, parallel (Fig. S10).

The supervised machine-learning classifier produced a lower classification accuracy than the rules-based classifier under leave-one-out cross-validation (0.70 vs 0.88; Supplementary Material, Section S2.1; Fig. S11).

### Circling and slinky swim

Circling was observed in seven shark tracks for durations of 7–45 min and was associated with turn radii of 4–10 m and rotation rates of 0.21–1.0 circles min⁻¹ (Fig. 3a–d). Compared with all other visually classified tracks >5 min, circling tracks (n=7) had reduced ground speed (0.58 ± 0.05 m s⁻¹ vs. 1.07 ± 0.26 m s⁻¹; Welch’s t(25.0) = −18.9, Bonferroni-corrected *p* = 1.5 × 10⁻¹⁵) and circling tracks meeting minimum tailbeat-data requirements (n=4) had lower tailbeat frequency (0.33 ± 0.02 Hz vs. 0.43 ± 0.09 Hz; Welch’s t(12.9) = −7.0, Bonferroni-corrected *p* = 5.8 × 10⁻^5^).

Slinky swim was observed five times and was associated with turn radii of 2.5–10 m and rotation rates of 0.21–0.42 circles min⁻¹ (Fig. 3e–h). Compared with all other visually classified tracks >5 min, slinky-swim tracks (n = 5) also had reduced ground speed (0.65 ± 0.06 m s⁻¹ vs. 1.06 ± 0.27 m s⁻¹; Welch’s t(8.2) = −12.1, Bonferroni-corrected *p* = 1.0 × 10⁻^5^) and those meeting the minimum tailbeat-data requirement (n = 4) had lower tailbeat frequency (0.29 ± 0.04 Hz vs. 0.43 ± 0.09 Hz; Welch’s t(4.6) = −6.2, Bonferroni-corrected *p* = 0.01) compared with all other visually classified tracks >5 min.

Among the broader movement modes, straight, parallel movement had higher ground speed than all other modes combined (1.13 ± 0.25 m s⁻¹ vs. 0.99 ± 0.27 m s⁻¹; Welch’s t(207.8) = 3.9, Bonferroni-corrected *p* = 7.2 × 10⁻^4^), but no other pairwise differences reached statistical significance (Tables S3 and S4).

### Distances from shore and directions of travel

Distance from shore and direction of travel (absolute and relative to shoreline) were calculated for 347 of the 359 processed tracks (shoreline data unavailable for some tracks). Measured distances ranged from 3–882 m (mean = 145 m; median = 116 m). Observation times increased linearly with distance out to 125 m (y = 0.06*x* − 0.87, R² = 0.93), then decreased (Fig. S12).

Average direction of travel across all tracks was primarily in the general north/south direction. When computed relative to shore using the boundary-aligned coordinate system, direction of travel distributions were tightened and shifted to be centered around the shoreline directions (Fig. 5a–b).

**Figure 5.**
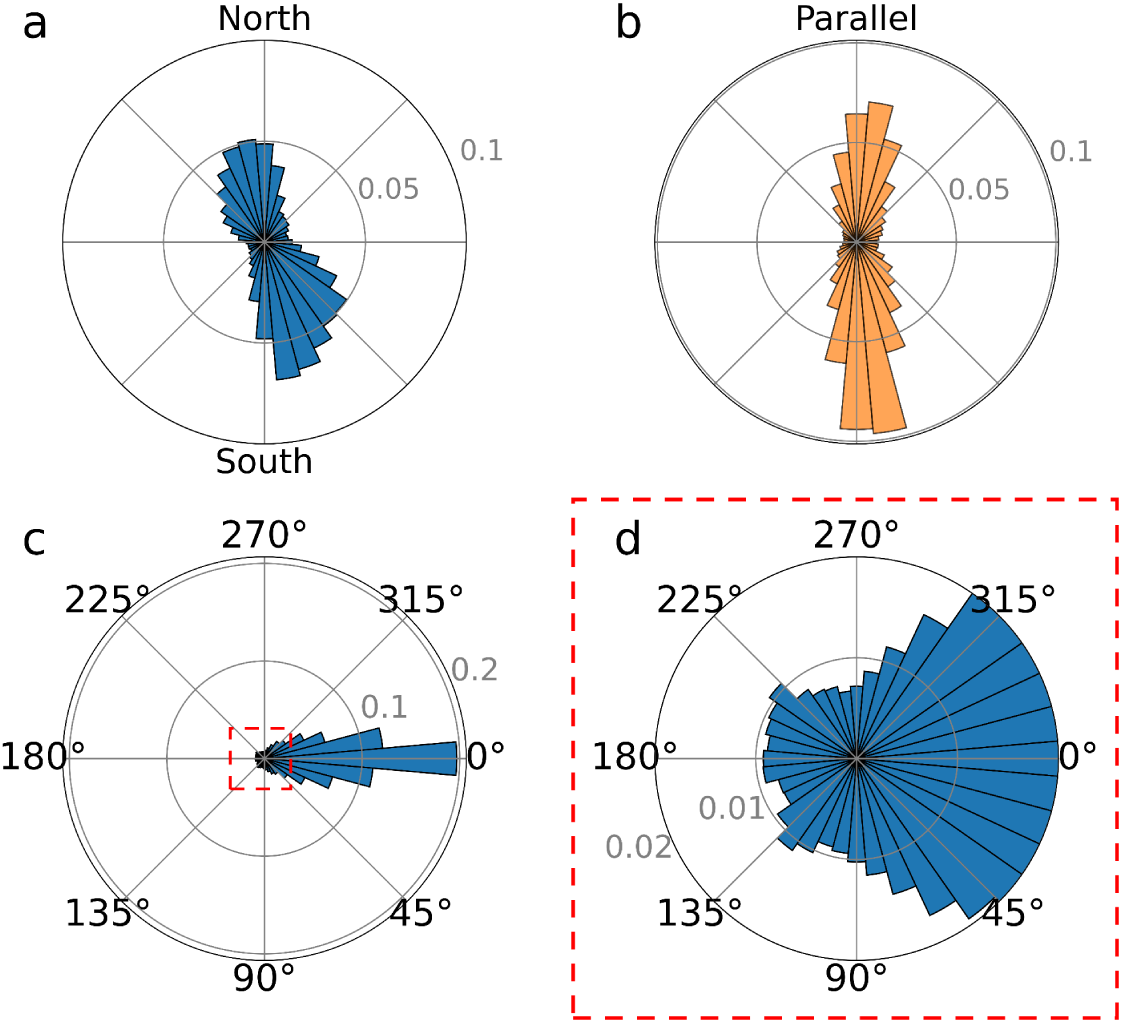
Polar histograms showing the distribution of shark directions of travel with bin widths of 10° and the radial axis representing the fraction of the total distribution in each bin. (a) Direction of travel shown as true heading. (b) Direction of travel shown relative to the shoreline where the distribution shifts and tightens around directions approximately parallel to the shoreline. (c) Direction of travel relative to the primary heading across 5-min track segments. The primary heading was defined as the most frequent bin in the distribution of instantaneous headings and was set to 0° for plotting. (d) Radial-scale zoom of the central portion of panel c, indicated by the dashed red box, with the maximum displayed bin radius limited to 0.02 to show lower-frequency directions. The concentration of headings near 0° indicates strong forward persistence over the 5-min segment durations examined; 62% of pooled heading observations were within ±30° of the primary heading, and 16% differed from the primary heading by more than 90°.

Sharks traveled approximately parallel (within ±30°) to shore 66% of the time, which is double the expected value of a uniform distribution. Across pooled 5-min segments, 62% of headings stayed within ±30° of the primary direction (Fig. 5c-d).

### Comparisons across beach locations

Descriptive differences in movement-mode distributions were observed among beach locations (Fig. S13, Table S5). Among non-transition tracks, Nauset exhibited the highest proportion of maneuvering, non-parallel tracks (33%, compared with 27% at Head of the Meadow and 14% at LeCount Hollow), although differences in parallel versus maneuvering, non-parallel mode counts did not reach statistical significance. Across all available track-level metric summaries meeting segment-analysis requirements, Nauset displayed values more consistent with non-parallel movement than the other beaches. NBAI, PBAI and BTI were all significantly lower at Nauset than at LeCount Hollow (NBAI: 0.85 vs. 0.96, Bonferroni-corrected *p* = 0.002; PBAI: 0.80 vs. 0.88, Bonferroni-corrected *p* = 0.003; BTI: 0.70 vs. 0.83, Bonferroni-corrected *p* = 0.012). NBAI and PBAI were also significantly lower at Nauset than at Head of the Meadow (NBAI: 0.85 vs. 0.97, Bonferroni-corrected *p* = 0.007; PBAI: 0.80 vs. 0.87, Bonferroni-corrected *p* = 0.041)

The distribution of heading directions relative to shore was also more dispersed at Nauset (Fig. S14). On average, tracks at Nauset spent 57% of valid heading time within ±30° of either shoreline-parallel direction, compared with 69% at Head of the Meadow and 75% at LeCount Hollow. Pairwise comparisons indicated that this difference was significant only between Nauset and LeCount Hollow (Dunn’s *p* = 8.9 × 10⁻^4^, Cliff’s δ = −0.41; Table S6).

## Discussion

We demonstrate that nearshore white shark movement is best understood not simply as movement through geographic space, but as movement relative to a persistent environmental boundary. Utilizing drone-derived trajectories captured off Cape Cod and a shoreline-referenced coordinate framework, we found that tracks were organized into recurrent boundary-related movement modes, dominated by sustained alongshore travel with localized maneuvering frequently embedded within broader shoreline-aligned movement. This structure indicates that the shoreline, and the bathymetric and ecological gradients associated with it, may act as a search corridor that organizes fine-scale predator movement. Several boundary-referenced metrics distinguished spatial patterns demonstrating that incorporating environmental geometry can reveal biologically meaningful structure. The approach provides both new insight into white shark nearshore behavior off of Cape Cod and a general framework for analyzing animal movement along persistent habitat edges.

### Biological context and observation constraints

Shark length measurements were consistent with previous studies from this sampling region (Supplementary Material, S3.1, Fig. S5) [20]. More than three quarters of track-level length measurements exceeded 3.0 m in total length, indicating that the sampled sharks were primarily composed of larger juvenile, subadult, and potentially adult individuals based on regional life-stage assessments [21,32]. These size classes are associated with coastal residency and an ontogenetic shift toward pinniped foraging in the western North Atlantic [27,33–35], linking movement patterns to the shoreline boundary and making this prey-rich coastal system well suited for examining fine-scale movement in a boundary-structured environment.

Ground speeds measured during nearshore tracking were typically on the order of 1 m s⁻¹, comparable to values reported previously for white sharks determined using both aerial methods and animal-borne sensors [12,36] and generally below estimates of the minimum cost of transport (≈1.3–1.9 m s⁻¹) [37]. While speeds are consistent with foraging, they may simply reflect energy-efficient localized movement and not necessarily active hunting [37,38]. Animals may traverse nearshore areas despite higher energy costs [37] so as to take advantage of potential scavenging opportunities or higher water temperatures [37,39]. Increased time in areas of high pinniped concentration may also afford animals the selectivity to engage only in the most opportunistic foraging opportunities.

Ground speeds scaled predictably with the product of tailbeat frequency and body length, consistent with established biomechanical relationships between swimming kinematics and translational velocity in fishes [40–42] (Supplementary Material, S3.3). Tailbeat frequencies fell within previously reported ranges [36,37,43]. However, they were consistently lower during localized circling and looping behaviors than during all other movements, indicating behavioral modulation of swimming kinematics associated with specific movement patterns which may reflect specific behaviors [44,45].

Drone-based data collection is inherently biased toward conditions where sharks are most visible from the air, complicating interpretation. As sightability generally decreases with increasing distance from shore, interpreting spatial distributions referenced to the shoreline must be done with caution. While observation time decreases beyond 125 m from shore, it is unclear to what extent this reflects reduced sightability as opposed to actual spatial distributions. Regardless, the linear increase in observation time with distance from shore out to 125 m (Fig. S12) suggests an actual increase in shark presence with distance from shore within this range. However, it must be noted that sandbar structure can disrupt the general relationship between distance from shore and water depth or sightability. These patterns further indicate that although detectability likely constrains offshore observations, sharks were not concentrated exclusively in the shallowest nearshore waters and were frequently encountered at moderate distances from the shoreline.

### Boundary-referenced metrics and movement modes

High-resolution movement data enable quantitative examination of fine-scale path geometry over ecologically relevant time scales. However, the interpretation of commonly used metrics is critically dependent on how they are calculated as well as the spatial context in which animals move. While metrics such as straightness provide intuitive measures of path tortuosity, their meaning is strongly scale-dependent and can be obscured when applied to tracks of variable length or intermittent continuity [18,19].

To quantify nearshore movement along a persistent linear boundary, we combined a number of scale-consistent metrics which included measures of straightness as well as a set of complementary metrics quantifying boundary alignment and another boundary-referenced descriptor quantifying nearshore confinement.

Shoreline alignment was captured by the NBAI which quantifies net displacement along the boundary and PBAI which is based on the cumulative orientation of movement along the boundary. PBAI captures the degree to which movement remains continuously aligned with a boundary. In systems where animals closely follow highly sinuous boundaries (e.g., rivers, channels, or reef edges), path-based measures such as PBAI provide an informative description of boundary-following behavior. As NBAI captures the overall directional progression along the boundary it is more effective at identifying sustained alongshore movement in situations where the path also oscillates despite overall movement along the shoreline (Fig. 2). As we defined these types of tracks as parallel in our visual assessment, NBAI provided an additional descriptor that was well suited to helping correctly recover visually assigned modes.

It is important to note that in the present system, the mapped shoreline likely represents only an approximation of the environmental boundary influencing shark movement. Animals may orient primarily with respect to nearshore bathymetry and associated ecological gradients, which generally run parallel to the shoreline but are not necessarily identical to it. Consequently, trajectories that follow the underlying environmental boundary may not remain perfectly aligned with the mapped shoreline.

Together, straightness, NBAI, PBAI and BTI provide an interpretable and parsimonious feature set that formalizes visually distinct movement patterns while avoiding reliance on track-level summaries or requiring latent state-inference models. This approach allows movement modes to be distinguished based in part on how trajectories are oriented and constrained relative to the shoreline, rather than solely on speed or turning rate.

Analysis of nearshore trajectories revealed a small number of recurrent movement patterns that were readily distinguishable by their geometry relative to the shoreline. Most tracks exhibited sustained boundary-parallel movement, either as relatively straight travel or as a maneuvering path of which the general direction remained aligned with the shoreline. Other tracks were characterized by spatially confined use of nearshore areas with frequent turning and circling behaviors. These patterns were visually apparent in raw trajectories and were consistently recovered using the boundary-referenced framework, supporting the interpretation that they represent distinct movement modes rather than continuous variation in path geometry.

Movement-mode composition varied with minimum track duration, with longer tracks more frequently capturing spatially confined, non-parallel movement. This effect likely reflects the greater temporal opportunity to observe localized maneuvering rather than a true absence of such behavior in shorter tracks (Fig. 4g), supporting the use of a 15-min threshold for summarizing movement-mode composition. Conversely, movements linking nearshore and offshore regions were less frequently observed over longer durations, consistent with reduced visual detectability as animals move offshore. These time- and visibility-dependent constraints highlight the importance of interpreting movement mode distributions in the context of observation duration and spatial context, particularly when using intermittent or spatially biased datasets.

As these limitations may lead to undercounting of nearshore–offshore transitions, we evaluated whether potential unobserved transitions were likely to bias interpretation of nearshore movement. When we identified additional potential transitions based on a less restrictive interpretation of track geometry and included observed dives, the overall pattern still indicated sustained inshore use relative to inferred transition frequency. This interpretation was further supported by the capture of multiple tracks on the order of hours (Supplementary Material, Note S4).

### Local variation in shoreline-associated movement

While differences in the prevalence of movement modes and path-based metric values among beach locations were generally modest, consistent trends suggest that local environmental context influences nearshore movement. Sharks observed at Nauset Beach exhibited greater directional variability, a higher proportion of non-parallel movement, and metric values more closely associated with maneuvering, non-parallel movement than at other beaches, particularly LeCount Hollow. This may reflect differences in bathymetry, sandbar complexity, or proximity to pinniped haul-out sites (Fig. 1) [24]. Similar associations between coastal geomorphology, prey distribution, and fine-scale predator movement have been reported at other nearshore sites, suggesting that boundary-structured movement may be shaped by local ecological conditions [46,47]. Interestingly, straightness values between the three locations did not show any statistically significant differences whereas a number of boundary-referenced metrics did. These included values for NBAI, PBAI and BTI between LeCount Hollow and Nauset (Table 1), suggesting these metrics may be better suited to identifying differences in movements that are shaped by the geography of the shoreline than are standard descriptors such as path tortuosity.

**Table 1.** Pairwise comparisons of track-level movement metric summaries across beach locations. Each row compares a movement metric (straightness, net boundary alignment index (NBAI), path boundary alignment index (PBAI), or boundary traversal index (BTI)) between two beaches. Values in parentheses indicate mean metric values calculated from mode-separated track-level summaries for each beach. Kruskal-Wallis tests were used to evaluate overall differences among locations, and Bonferroni-corrected p-values from post hoc Dunn’s tests are reported alongside Cliff’s δ effect sizes and qualitative effect magnitudes. Most comparisons showed small or negligible effects; however, medium-sized differences were observed between Nauset and LeCount Hollow for all three boundary-referenced metrics, and between Nauset and Head of the Meadow for NBAI and PBAI. Sample sizes were identical across all metrics within each beach: Head of the Meadow, *n* = 39; LeCount Hollow, *n* = 52; and Nauset, *n* = 40 mode-separated tracks. Site abbreviations are used for compactness: HM = Head of the Meadow and LeCounts = LeCount Hollow.

| Metric | Beach 1 (value) | Beach 2 (value) | Group Test (Kruskal-Wallis) | Bonferroni-corrected p-value | Effect Size (Cliff's $\delta$) | Effect Magnitude |
| --- | --- | --- | --- | --- | --- | --- |
| **Straightness** | LeCounts (0.85) | Nauset (0.76) | H = 3.52, $p = 0.172$ | 0.201 | 0.229 | Small |
|  | HM (0.76) | LeCounts (0.85) |  | 0.745 | -0.136 | Negligible |
|  | Nauset (0.76) | HM (0.76) |  | 1.000 | -0.074 | Negligible |
| **NBAI** | LeCounts (0.96) | Nauset (0.85) | H = 13.67, $p = 0.001$ | 0.002 | 0.399 | Medium |
|  | HM (0.97) | LeCounts (0.96) |  | 1.000 | -0.036 | Negligible |
|  | Nauset (0.85) | HM (0.97) |  | 0.007 | -0.415 | Medium |
| **PBAI** | LeCounts (0.88) | Nauset (0.80) | H = 11.57, $p = 0.003$ | 0.003 | 0.403 | Medium |
|  | HM (0.87) | LeCounts (0.88) |  | 1.000 | -0.079 | Negligible |
|  | Nauset (0.80) | HM (0.87) |  | 0.041 | -0.319 | Small |
| **BTI** | LeCounts (0.83) | Nauset (0.70) | H = 8.63, $p = 0.013$ | 0.012 | 0.368 | Medium |
|  | HM (0.78) | LeCounts (0.83) |  | 1.000 | -0.067 | Negligible |
|  | Nauset (0.70) | HM (0.78) |  | 0.127 | -0.241 | Small |

### Boundary-aligned movement as a dominant nearshore strategy

Shoreline-parallel movement by white sharks has been documented previously [12,46,47], indicating that boundary-parallel movement is not simply an artifact of aerial sampling but a recurrent feature of coastal behavior. Our results extend this work by demonstrating that this alignment is not merely incidental but represents a dominant and temporally persistent geometric mode of nearshore space use. Some individual tracks demonstrated persistent boundary-aligned movement for extended periods (up to several hours), suggesting that the movement modes identified here can persist over time scales substantially longer than typical observation windows.

Directional headings were concentrated within ±30° of the shoreline approximately twice as frequently as expected under isotropic movement, indicating structured boundary-oriented travel rather than random displacement constrained by geography alone. Importantly, maneuvering behavior did not necessarily disrupt this alignment; instead, oscillatory or looping movements were often embedded within an overall shoreline-aligned trajectory. This pattern suggests that the coastline functions as a structured search corridor, reducing effective search dimensionality while allowing localized inspection of nearshore areas.

Although parallel movement along shorelines may appear intuitive given the linear geometry of coastlines, several alternative movement strategies would have been plausible, including localized patch residency near prey aggregations, frequent movement between offshore and nearshore habitats, purely linear transit or largely isotropic movement constrained only by bathymetry. The dominance of sustained boundary-aligned trajectories observed here suggests that nearshore movement is structured primarily as extended interface-following search rather than discrete patch use or repeated habitat transitions.

### Broader implications for boundary-referenced movement analysis

The agreement between visually identified modes and those classified quantitatively further supports the use of boundary-referenced metrics to formalize movement patterns that are readily recognized by observers but are otherwise difficult to compare across studies. Persistent linear boundaries are common organizing features across ecological systems, including reef margins, riverbanks, ice edges, and anthropogenic habitat interfaces. In such environments, movement relative to boundary geometry may shape encounter rates, sensory information flow, and energetic expenditure. By explicitly incorporating environmental structure into trajectory analysis [48], boundary-referenced approaches provide a transparent means of resolving recurrent movement modes in spatially constrained habitats. The framework developed here is therefore not specific to white sharks or coastal systems but may be broadly applicable to animals operating along persistent habitat edges.

## Data Availability Statement

All processed data reported in this study along with the code used to generate figures and conduct the analyses detailed here will be made available via Code Ocean upon peer-reviewed publication.

## Acknowledgements

The authors would like to thank Mark McCormick, David Sexton, Taylor Lamme, David Mizrahi, Malcolm Stewart, Josh Stewart, Geoff Reuland, John Reuland and Jackson R. Davis for their assistance with field experiments and piloting. They also wish to thank Simon Benhamou, who provided valuable insight as we prepared this manuscript. The authors would like to further acknowledge support from the Kay Fellowship through the ‘Iolani School. Finally, they would like to thank the Cape Cod Ocean Community and the Dorr Foundation for their support.

