## Supplementary Material for "A boundary-referenced framework for quantifying shoreline-associated movement in white sharks"

**Author Affiliations**
1. Moosh Systems, LLC. Wellfleet, MA, USA
2. The Cooper Union for the Advancement of Science and Art, New York, NY, USA
3. Biology Department, Stanford University, Stanford, CA, USA

* Corresponding Author

### Supplementary Material

#### Note S1. Supplementary Methods

This supplementary methods section provides detailed descriptions of data collection procedures, annotation protocols, calculation methods, and filtering criteria used in the collection and analysis of drone-based shark tracking data presented in the main manuscript. These methods support and expand on the summary provided in the primary Methods section, and are included here to facilitate reproducibility and methodological transparency.

##### S1.1 Video and Flight Record Alignment and Processing

All video was captured using onboard microSD cards at 4K resolutions of either 3840x2160 or 4096x2160 pixels and frame rates of either 30 or 60 frames per second (fps). The aircraft allows for capture of flight records at up to 10 Hz, which include altitude, magnetic heading, gimbal angle, and GPS coordinates. Custom Python scripts were used to align and process all video data and flight records as detailed in Sexton et al. [1].

##### S1.2 Shark Lengths, Ground Speeds, and Tailbeats

Shark lengths and kinematic metrics were extracted from annotated video segments under specific visibility and geometry criteria using Python scripts, as outlined below.

Shark length measurements were taken when the animal was at or near the surface (estimated depth < 1.5 m) with its outline clear and the drone at altitudes between 46-122 m with the gimbal at or above 50°. During the annotation process the user was able to select optimal images with minimal distortion on which to take measurements irrespective of the animal's tail position. To account for variable body positions while swimming the user selected a series of points along the centerline of the shark and distances between all points were calculated and summed. This allowed the user to take multiple measurements as well as select the clearest images regardless of body position. In cases where multiple clear annotated images were available for the same shark, reported lengths are averages from all images [1].

Shark speeds were calculated from filtered position and time data for each individual track. The resulting raw speed estimates were then filtered using a centered 30-s rolling median absolute deviation filter to remove spurious high-speed values arising from localized positional artifacts; values exceeding 4.0 m $s^{-1}$ were also excluded as an absolute artifact threshold and finally values associated with recorded tailbeat frequencies ≥ 2.0 Hz were discarded.

Shark tailbeat frequencies were recorded manually during the annotation process by marking the times where the tail reached full lateral deflection in each direction. This procedure was performed only when tail reversals could be resolved unambiguously. All sequences included in analysis had tailbeat frequency measured for at least 0.5 minutes. All reported tailbeats are averages over the duration of measurements for the entirety of each track with any periods of burst swimming as defined by tailbeats ≥ 2.0 Hz discarded [2].

**S1.3 Shoreline Mapping**

As shorelines along the outer Cape shift on a regular basis, accurate maps are not readily available. Horizontal shark position and direction of travel relative to the shoreline were determined by mapping the coastline using the drone. Shorelines were mapped by flying the drone directly over the shoreline with the gimbal at the nadir.

Custom Python scripts allowed the user to pause videos and draw a series of points over the shoreline using a touchscreen (Fig. S2). Shorelines were drawn intermittently throughout shoreline-dedicated videos such that there was generally some overlap in marked points. Points were smoothed and converted to geographic coordinates using the same alignment and projection techniques used for shark position data. In instances where there was no overlap of shoreline markings, linear interpolation was used to fill any shoreline gaps less than 100 m. Shoreline coordinates were then smoothed using a moving mean filter and cubic interpolation was used to establish coordinates at 0.3 m resolution.

Shark and shoreline coordinates were transformed into a common local planar coordinate system. The ordered shoreline coordinates were further smoothed using a centered 200 m moving mean filter and then connected by straight line segments to form a continuous shoreline polyline. For each filtered shark position, the nearest point anywhere along this polyline was first calculated as a target. To reduce abrupt shifts caused by local ambiguity in nearest-point associations, movement of the accepted shoreline point was step-limited within continuous shark-track sections. The first valid position in each section was assigned directly to its nearest shoreline point. For each subsequent position, the accepted shoreline point was allowed to move in either direction along the polyline toward the nearest-point target by no more than twice the shark's displacement between successive filtered positions. The accepted association therefore equaled the nearest-point target when it fell within this limit but otherwise remained between the previous accepted association and the new target. This retained genuine reversals in alongshore movement while constraining disproportionately large changes in shoreline association. The constraint was reset at track discontinuities.

Accepted shoreline associations were used to calculate each animal's distance from shore over the portion of its track for which shoreline positions were available. All distances were corrected for tide changes between the time of shoreline acquisition and the time of each shark position using a polynomial fit correlating tide height with shoreline change.

Shoreline and shark positional accuracies were estimated based on the log-normal error distributions established in Sexton et al. (11.4 m; 95% CI) [1]. The error in distance to shore measurements was calculated using these error estimates in conjunction with the root-sum-of-squares method resulting in an estimated error of 16.1 m (95% CI) [1,3]. For tracks where both the shark and shoreline were at times seen within the same field of view, direct measurements were used to calibrate distances to shore calculations across those tracks [1].

##### S1.3.1 Local Shoreline Bearing

Local shoreline bearing was estimated at each accepted shoreline location to approximate the local tangent direction. The corresponding position on the shoreline polyline was identified, and shoreline bearing was calculated from adjacent shoreline coordinates using a centered-difference approach. Tangent vectors were normalized and checked for directional continuity. Because the eastern Cape Cod shoreline is approximately linear over the spatial scales analyzed, this local tangent approximation provides a stable estimate of shoreline orientation. All angles were computed using the two-argument inverse tangent function (atan2) to preserve full-quadrant directional information.

#### S1.4 Shark Track Grouping and Filtering

After obtaining geographic positions and mapped shorelines, shark movements were grouped into tracks for further analysis. All instances in which visual contact was lost required assessment of the likelihood that a subsequent track represented the same shark. In most cases this assessment was performed by examining plots of shark latitude versus time. If linear fits across gaps in these plots did not match the slope of the trajectory on either side of the gap, the tracks were considered to represent different sharks.

On the few occasions in which sharks were observed to be continually maneuvering in one small area, tracks were considered to be of the same shark if the tracks exhibited similar maneuvering patterns in that same location. Additionally, whenever possible, shark length measurements were made with a 0.5 m mean difference in length used as the cutoff to classify the animals as the same or different. In no cases in which sharks were initially categorized as the same individual did length measurements exceed this threshold and require the reclassification of animals. These procedures were used to determine whether observations separated by temporary losses of visual contact belonged to the same continuous tracking encounter. Individuals could not generally be reidentified across tracking encounters conducted on different dates or at different locations; tracks were therefore treated as observational units, and the same shark may have contributed more than one track.

If at any time during data collection the animal being tracked was approached by a research vessel, the track from that point on was discarded.

##### S1.5 Segment-Based Analysis

Tracks were divided into contiguous segments by splitting at temporal gaps exceeding 30 s. Segments shorter than 5 min were excluded. Within each segment, overlapping 300 s windows were defined at 15 s intervals, and metrics were calculated independently for each window.

To obtain point-wise values, each window-level metric was assigned to all positions within that window. Because windows overlapped, individual positions were typically associated with multiple windows; final values were therefore calculated as the mean of all window-level values corresponding to that position.

This procedure produces a continuous representation of movement properties along tracks equivalent to a centered moving-window calculation while avoiding the use of truncated windows.

##### S1.6 Rules-Based Classification Cross-Validation

Thresholds defining the rules-based classifier were optimized using coordinate descent to maximize per-track classification accuracy on the training data. To assess generalization performance, we implemented leave-one-out cross-validation (LOOCV). For each of the 131 tracks, a single track was removed, thresholds were re-optimized on the remaining 130 tracks, and the held-out track was classified using the refitted thresholds. Accuracy was recorded for each fold. Because thresholds were re-estimated at each iteration, LOOCV provides an estimate of predictive performance for this modest-sized dataset.

Robustness was further evaluated using nonparametric bootstrap resampling (B = 1000 replicates). For each replicate, 131 tracks were sampled with replacement, thresholds were optimized on the bootstrap sample, and classification accuracy was evaluated on out-of-bag (OOB) tracks not included in that sample. All replicates produced non-empty OOB sets. The distribution of OOB accuracies was used to estimate predictive performance variability.

##### S1.7 Machine Learning Classification

To complement the rule-based classification approach, we implemented a supervised machine learning model using a Random Forest classifier from the scikit-learn Python package. The random forest contained 100 decision trees and was fitted without class weighting or additional hyperparameter tuning. The model was trained on point-level metric records from visually assigned, mode-separated tracks using straightness, net boundary alignment index (NBAI), path boundary alignment index (PBAI), and boundary traversal index (BTI) as inputs. For training, multi-modal initial tracks were separated into single-mode analytical tracks.

Model predictions were then applied to point-level metric records from the combined, unseparated track dataset. Predictions were summarized for the corresponding mode-separated analytical tracks, with the overall mode assigned as the most frequent predicted mode. Leave-one-out cross-validation was conducted across the 131 mode-separated tracks meeting continuity requirements. To avoid leakage among modes derived from the same initial track, all records from the same parent shark ID were excluded from the training set when one of its mode-separated tracks was held out.

#### Note S2. Supplementary Results

##### S2.1 Machine Learning Classification

Machine learning classification using leave-one-out cross-validation demonstrated lower performance than the rule-based approach. Across 131 tracks, accuracy was 67% at the point level and 70% when tracks were assigned their most frequent mode (Fig. S11). Most errors occurred between straight, parallel and maneuvering, parallel.

Although performance was lower than that of the rule-based classifier, the Random Forest results indicate that the four movement metrics captured substantial structure separating behavioral modes. Given the modest sample size and low-dimensional feature space, the stronger performance and greater transparency of the rule-based system supported its use for the primary analyses.

#### Note S3. Supplementary Analyses and Discussion

##### S3.1 Length Measurements: Comparison to Previous Studies

This supplemental discussion section provides a more detailed comparison of the measured lengths reported here to those found in previously published datasets.

Traditionally, reported shark lengths have relied on visual judgement [4–9] or references within images [10,11]. While recent drone studies have reported lengths without references within images, these measurements have not been accompanied by detailed error analysis [12,13], and other studies have refrained from reporting lengths at all [14]. Low-cost methods to accurately measure lengths [1], such as those used here, are critical for characterizing population structures [10,15,16].

Shark lengths determined during the current study were compared to those reported by Winton et al. [11]. Length data from Winton et al. were extracted from published histograms to enable approximate comparison of distributional ranges and central tendencies. The two datasets show substantial overlap in total length distributions, with similar ranges and broadly comparable central tendencies (Fig. S5), indicating consistency in the size structure of sharks observed across studies despite differences in sampling period and measurement procedures.

Both studies utilize aerial imagery; however, given the differences in methods, survey platforms and sampling periods [1,11], we do not necessarily interpret small differences in summary statistics as evidence of biological change or systematic measurement bias. Rather, this comparison is intended to demonstrate that the size distributions observed in the present study fall within the range previously reported for nearshore white sharks in this region.

##### S3.2 Sampling Context and Limitations of Length Data

Length measurements reported here, as well as those reported by Winton et al. (2023), are derived from nearshore sampling along the eastern coast of Cape Cod and therefore characterize only the subset of the regional white shark population that utilizes this shallow coastal habitat [11]. Regional tagging and sampling data from nearby areas suggest that sampled size distributions can vary with habitat use and survey location [17]. Consequently, the length data presented here should be interpreted as representative of nearshore-resident sharks rather than the full size structure of the region surrounding Cape Cod.

##### S3.3 Speed Measurements

This supplemental section provides expanded comparisons between ground speeds measured in the current study and those reported in previous studies using drones and biologging tags. Differences in measurement techniques and the effects of environmental factors such as currents are also discussed.

Ground speeds measured in the present study ranged from 0.29 to 1.81 m $s^{-1}$, with a mean of 1.04 m $s^{-1}$ across 288 tracks (Fig. S4b). These values are comparable to ground speeds reported previously for white sharks using drone-based observations (0.54–1.23 m $s^{-1}$) [12] and to swim speeds inferred from animal-borne accelerometers (0.8–1.35 m $s^{-1}$) [18]. Average speeds were generally below published estimates of minimum cost of transport (≈1.3–1.9 m $s^{-1}$) [5].

Drone-derived speeds represent displacement over the ground and as such are susceptible to movement artifacts and also do not explicitly account for ocean currents, in contrast to swim speeds estimated from animal-borne sensors. Along the eastern shore of Cape Cod, longshore currents commonly range from approximately 0.1 to 0.2 m $s^{-1}$ and can be higher [19], which may contribute to variation in measured ground speeds. As a result, small differences in reported speeds across studies should be interpreted cautiously and not over-attributed to behavioral or physiological differences. Tailbeat frequency may in fact better reflect subtle differences in transport efficiency and energy use [20]. While tailbeat data were extracted using manual methods here, more advanced image processing will allow for more automated procedures [21].

Tailbeat frequencies measured in this study ranged from 0.24 to 0.63 Hz (mean: 0.43 ± 0.09 Hz; median: 0.43 Hz) and were consistent with values reported previously for white sharks engaged in nearshore and foraging-associated behaviors (Fig. S4c) [18,21]. Tailbeat frequency was lower during circling and looping behaviors than during all other behaviors, indicating behavioral modulation of swimming kinematics associated with specific movement patterns (Table S4).

Importantly, ground speed scaled linearly with the product of tailbeat frequency and body length across track-level observations, consistent with established biomechanical relationships between swimming kinematics and translational velocity in fishes (Fig. S4d) [2,20,22].

##### S3.4 Durations of continuous inshore activity

In order to understand the extent to which nearshore–offshore transitions may be undercounted, we categorized all tracks as either potential transition or non-transition. Tracks where animals were lost due to dives or as they moved offshore as well as tracks where sharks were initially found moving toward the shoreline were classified as potential transitions. This resulted in 48 of 359 tracks (13%) containing potential transitions.

We estimated the duration of continuous inshore activity by comparing total tracking time to instances of potential nearshore–offshore transitions. With ~68 hours of total tracking this translated to ~1.4 hours of inshore activity per potential transition (48 potential transitions). Here total tracking time is defined as the total time from the start of tracking to the end of tracking and includes time between periods of active tracking in which the animal position is unknown but the animal is assumed to remain inshore. As such, total tracking time is considerably higher than active tracking time which includes only time for which the animal position is known.

While we consider all active tracking to be inshore, tracking beyond even 300 m from shore is limited and so dropping tracking time for animals further offshore will have minimal effect on estimates (Fig. S12).

Our analysis assumes that transitions between nearshore and offshore environments are not more likely to occur under conditions of lower sightability. If animals were to preferentially transition at deeper swim depths then transitions could be significantly undercounted. While we have no evidence to suspect this is the case, we do not have data to rule out this possibility. However, the capture of multiple continuous tracks on the order of hours tends to support our estimates for continuous inshore activity (Note S4).

#### Note S4. Descriptions of Long-Duration Shark Tracks

The ability to capture long-duration tracks was limited during this study as a significant portion of flights were conducted from fixed locations on the beach and the vast majority of the time only a single drone was used. However, data collection in 2022 focused on attempts to capture long-duration continuous tracks and as such flights were conducted from a research vessel utilizing multiple drones. These efforts were limited to 7 research trips of between 4 and 8 hours in length with an estimated 23 shark sightings and 15 tracking events. During these trips, single shark tracks in excess of 1 and 3 hours were captured and are detailed here. An additional track is suspected to be of a single shark maneuvering in a limited area for nearly 3 hours but several instances where visual contact was lost in conjunction with the presence of other sharks in the area prevent us from definitively concluding the track is of a single shark. The capture of multiple tracks measured in hours during this limited sampling effort suggests that our estimated average continuous inshore movement duration of 1.4 h is reasonable and may in fact be an underestimate (Section S3.4).

##### S4.1 Track 1 – Wellfleet (September 2022)

A white shark was tracked continuously for 72 minutes, as it swam southward for 4.2 km along the Wellfleet shoreline from just south of Whitecrest Beach past LeCount Hollow all the way past Marconi Beach. The animal maintained a largely straight, parallel course relative to the shoreline, with a standard deviation in heading relative to shore of 17°. The shark remained within 72 to 196 m of the shoreline (mean = 140 m; SD = 32 m), passing multiple beaches and three groups of surfers. Tracking was discontinued upon depletion of available batteries with the shark still clearly visible and still headed along the same course.

##### S4.2 Track 2 – Nauset (September 2022)

A white shark was tracked nearly continuously for approximately 3 hours and 33 minutes (interrupted by a <10-minute visual loss after which the animal was reacquired and identified via distinctive markings). The shark remained within a 2.3 km section of coast near Nauset Beach, traveling repeatedly up and down the shoreline. The animal's distance from shore ranged from 9 to 441 m (mean = 197 m, SD = 94 m), and its movement was characterized by maneuvering behavior with frequent changes in direction.

### Supplementary Figures

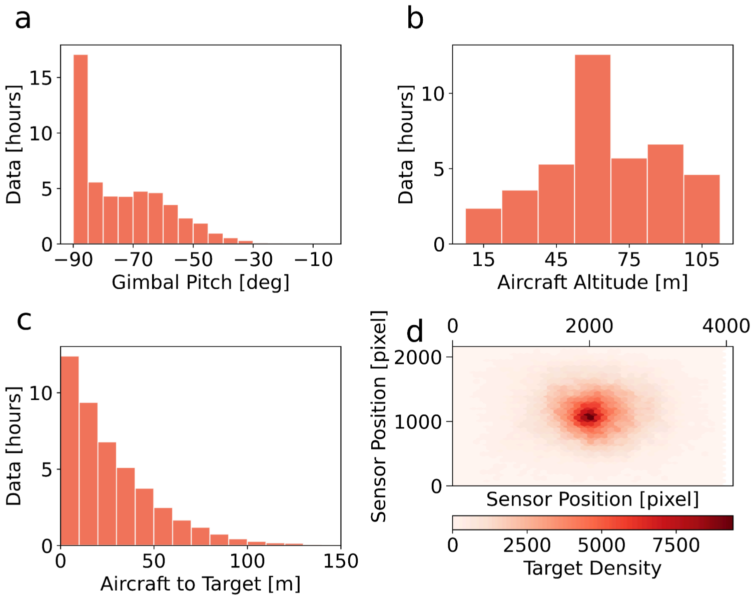


**Figure S1**. Data collection parameters over the course of the study in which 50.4 hours of tracking data were collected and 359 separate shark tracks were processed. (a) Gimbal angles. (b) Flight altitudes ranged from <15 m to 122 m. (c) Horizontal distances from aircraft to target determined by gimbal angle, altitude, and target position on the image sensor. (d) Density of target positions on the image sensor. Pixels are binned into hexagons, each covering 102 pixels horizontally and 52 pixels vertically.

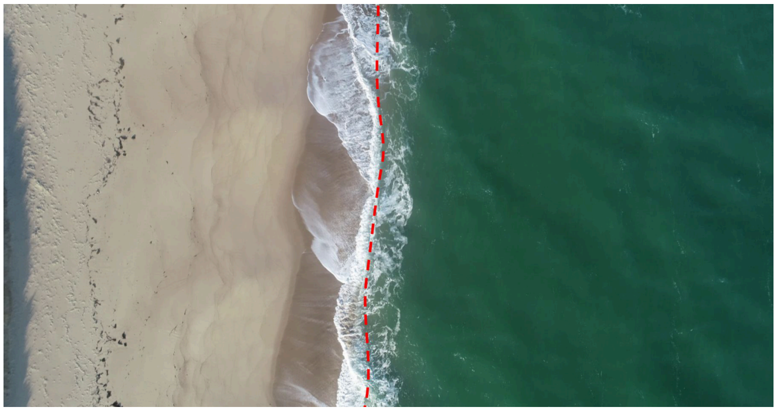

**Figure S2**. Example image showing shoreline mapping from drone video. The drone was flown along the shoreline with the gimbal at the nadir. Custom Python scripts enabled users to pause video playback and trace the shoreline using a touchscreen (red dashed line). Shoreline positions on the image sensor, combined with image and flight data, were used to create shoreline maps for calculating shark position and direction of travel relative to the shoreline.

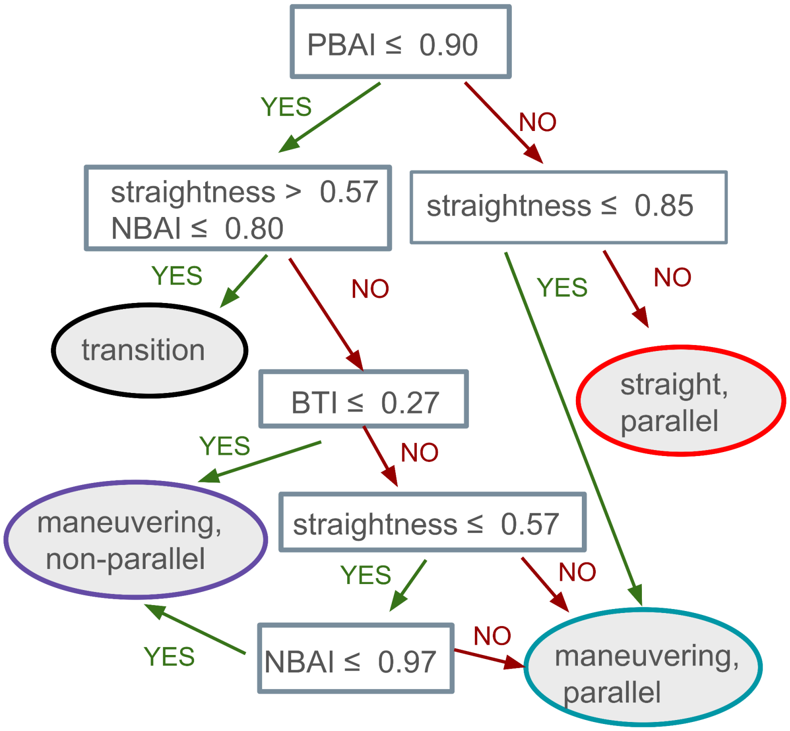


**Figure S3**. A rules-based classification system utilized a decision tree with six conditional if-then statements defining branches. The tree utilized four track-based metrics for classification: straightness, net boundary alignment index, path boundary alignment index and boundary traversal index. The rule structure reflects a hierarchical geometric interpretation of shark movement relative to the shoreline. Tracks were first separated using path-level shoreline alignment because this metric captures whether the shark's realized path generally followed the shoreline, independent of whether the movement was straight or tortuous. Tracks with high path alignment were therefore treated as shoreline-parallel and subsequently divided into straight versus maneuvering movement based entirely on straightness. In contrast, tracks with lower path alignment were treated as geometrically ambiguous and evaluated using additional criteria. Within this branch, relatively straight tracks with low net shoreline alignment were classified as transitions, consistent with directed movement across or away from the shoreline. Tracks with low boundary traversal were classified as maneuvering, non-parallel because they showed limited alongshore displacement. Remaining low-path-alignment tracks were then separated using straightness and net shoreline alignment, allowing irregular or tortuous tracks with strong net alongshore displacement to remain classified as maneuvering, parallel.

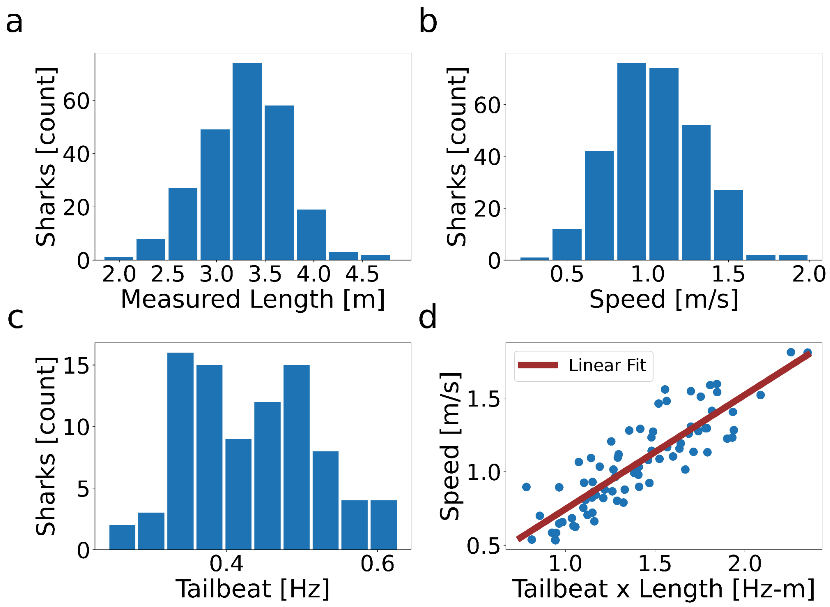


**Figure S4**. Summary of shark lengths, speeds, tailbeat frequencies, and their relationship to one another. (a) Length distribution from 241 shark tracks. Lengths ranged from 1.97 m to 4.63 m (mean ± SD: 3.30 ± 0.44 m; median: 3.35 m), with 77% exceeding 3.0 m. (b) Distribution of average ground speeds for 288 tracks, ranging from 0.29 to 1.81 m $s^{-1}$ (mean: 1.04 ± 0.27 m $s^{-1}$; median: 1.04 m $s^{-1}$). A Shapiro-Wilk test did not show evidence of non-normality (p = 0.54). (c) Distribution of average tailbeat frequencies for 88 tracks, ranging from 0.24 to 0.63 Hz (mean: 0.43 ± 0.09 Hz; median: 0.43 Hz). A Shapiro-Wilk test did not show evidence of non-normality (p = 0.09). (d) Ground speed plotted against the product of length and tailbeat frequency (n = 79). Linear regression yielded a slope of 0.77 (SE = 0.05), with an intercept of –0.03, $R^2$ = 0.87, and p = 7.3 × $10^{-25}$.

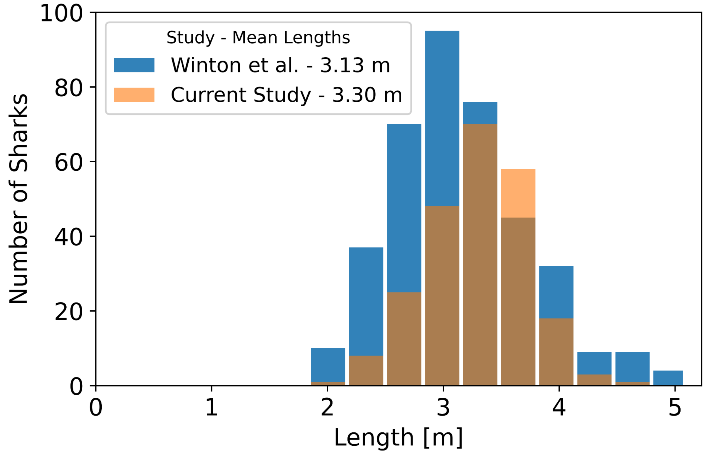


**Figure S5**. Histogram of shark length measurements from the current study overlaid on those reported by Winton et al. The mean total length in the current study was 3.30 m as compared to 3.13 m reported by Winton et al. The distributions share substantial overlap with similar ranges and shapes despite differences in sampling period and measurement procedures.

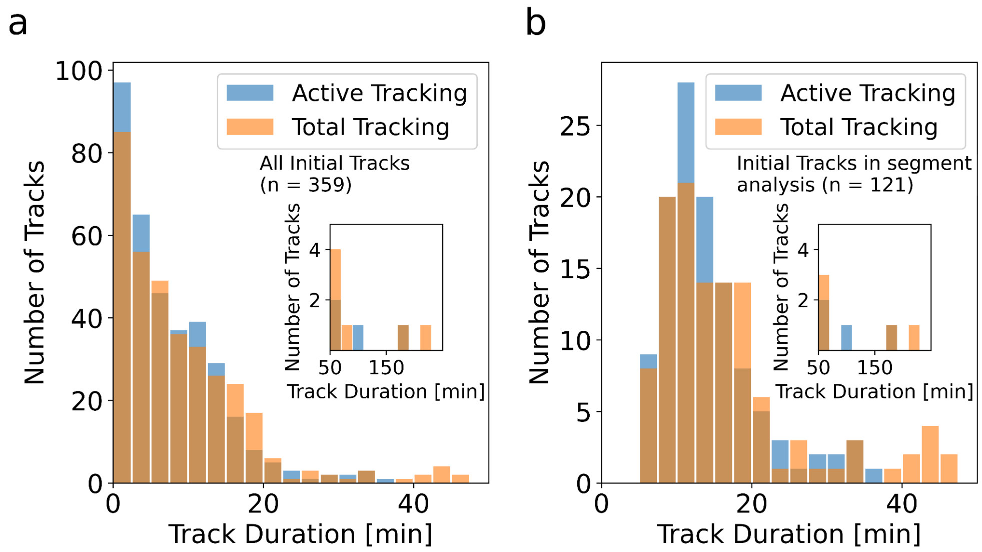


**Figure S6**. Histograms showing the duration of individual shark tracks. (a) Includes all initial tracks (n=359). (b) Includes only initial tracks meeting continuity requirements for segment analysis (n=121). Total tracking represents the time from the start of tracking to the end of tracking and may include gaps in which the shark position is unknown. Active tracking is the total time for which the shark position can be determined by the aircraft. Insets provide a view of longer track duration data not shown in main plots.

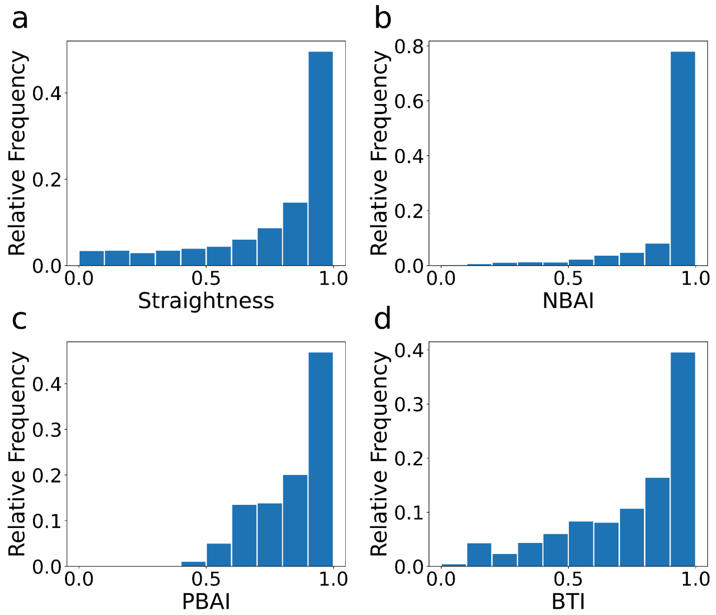


**Figure S7**. Histograms of straightness, net boundary alignment index, path boundary alignment index, and boundary traversal index calculated from overlap-averaged point/time-sample values derived from 5-min segment metrics. Mean values were 0.77 for straightness, 0.92 for NBAI, 0.84 for PBAI, and 0.75 for BTI.

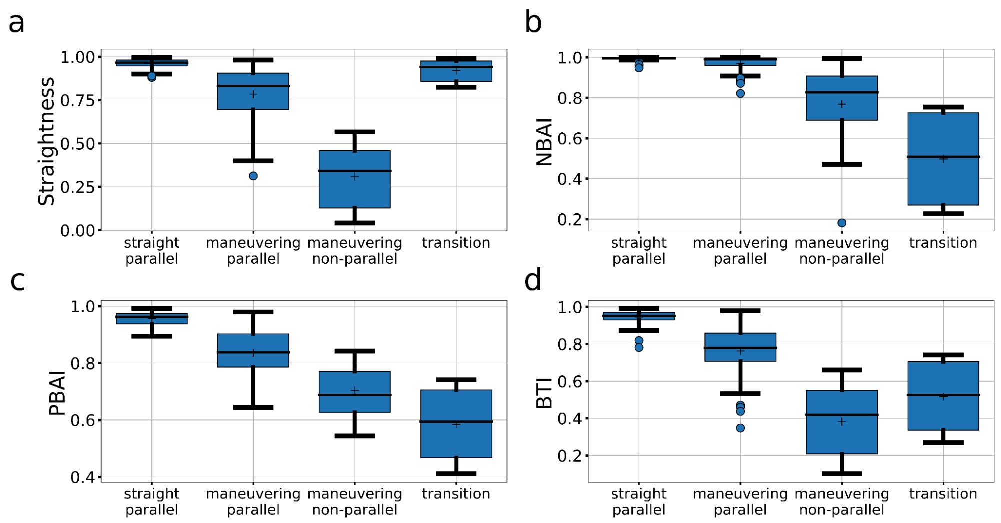


**Figure S8**. Boxplots of mode-separated track metric values across visually determined movement modes. Individual tracks were visually examined, with multimodal tracks split into single-mode analytical units. Of the resulting mode-separated tracks, 131 met the requirements for segment analysis. Mode sample sizes were straight, parallel: n = 52; maneuvering, parallel: n = 55; maneuvering, non-parallel: n = 18; and transition: n = 6. Calculated metrics included (a) straightness, (b) net boundary alignment index, (c) path boundary alignment index, and (d) boundary traversal index. Boxes show the interquartile range (IQR), whiskers extend to the most extreme data point within 1.5×IQR from the box, and outliers are plotted individually. Means are indicated by plus signs (+).

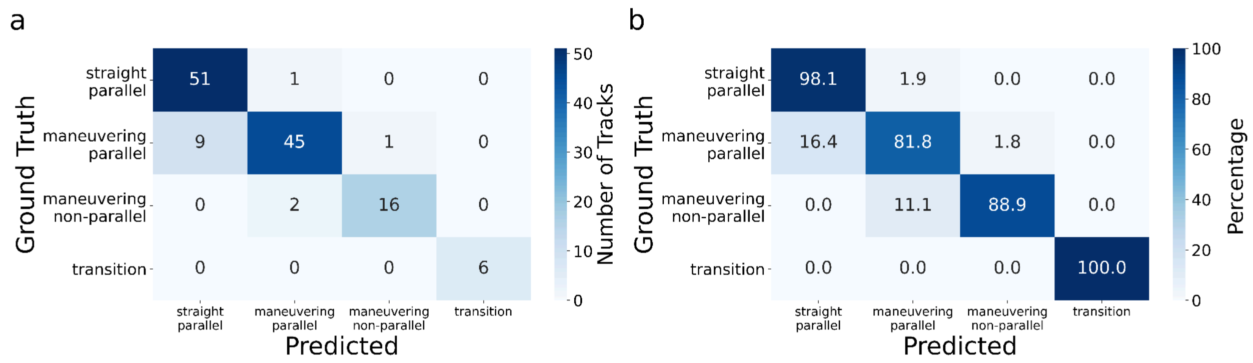


**Figure S9**. Confusion matrix showing the performance of the rule-based movement mode classification optimized using all 131 mode-separated tracks that independently met segment analysis requirements. Predicted modes based on average metrics were assigned to all tracks. Rows represent the ground truth (visual classification) and columns represent the predicted modes. (a) Each cell indicates the number of tracks with the corresponding true and predicted classification. (b) Each cell indicates the percentage of tracks in that row with the corresponding true and predicted classification. Overall agreement was 90% for the final threshold set optimized using all tracks (118/131) and balanced accuracy, calculated as the mean of class-specific recall values across movement modes, was 92%. For the leave-one-out cross-validation analysis, overall accuracy was 0.88 (115/131) and balanced accuracy 0.90.

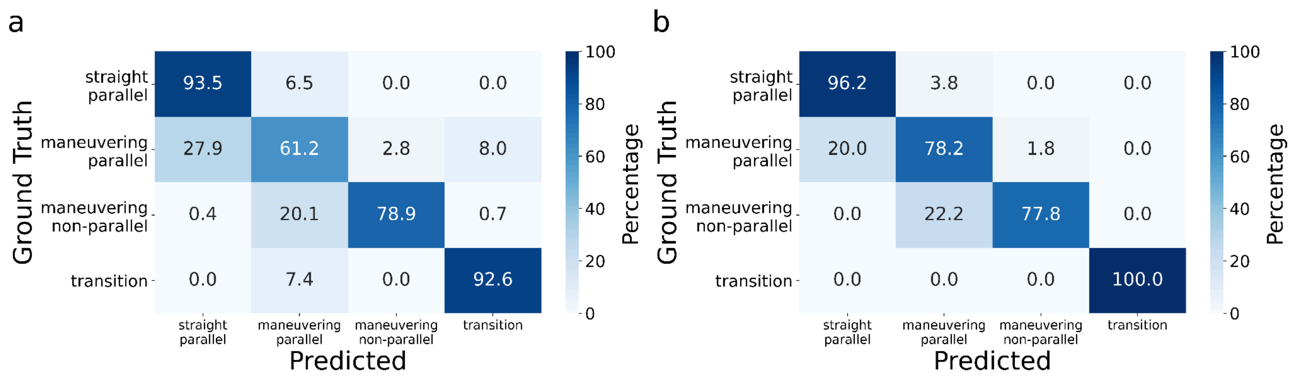


**Figure S10**. Confusion matrix showing classification performance of dynamic rule-based mode prediction for 131 mode-separated tracks meeting segment analysis requirements. All metrics used by the classifier, straightness, net boundary alignment index, path boundary alignment index, and boundary traversal index, were calculated from segments derived from full initial tracks prior to any visual separation of tracks into individual modes. Predictions were made independently for each point based on these dynamic metrics. For evaluation, points were assigned to visually assessed mode-separated tracks, and performance is shown both at the point level and after majority-vote summarization at the mode-separated track level. Overall agreement at the point level was 0.76 and at the track level 0.86.

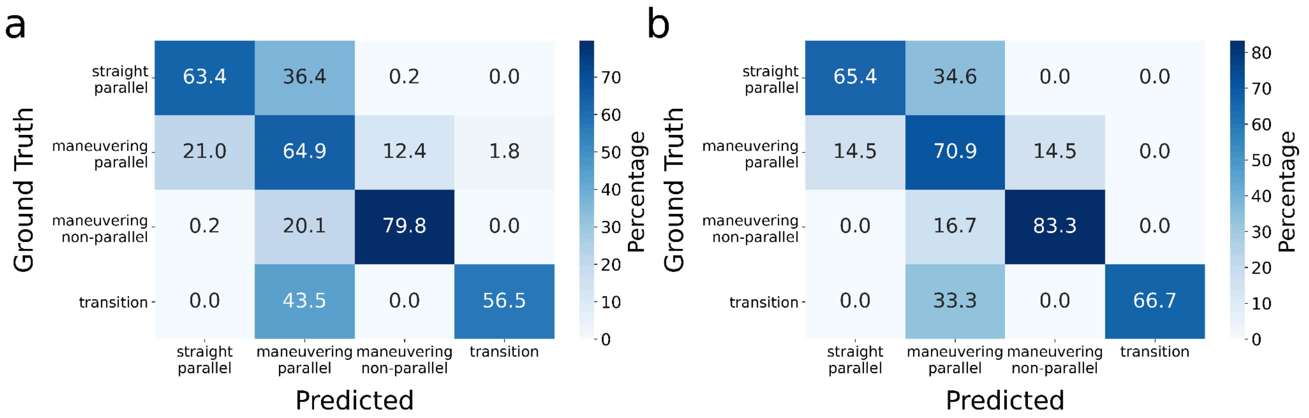


**Figure S11**. Confusion matrices showing classification performance of the Random Forest classifier for dynamic movement mode prediction. Rows represent visually assigned movement modes and columns represent predicted modes. Values are row-normalized percentages, indicating the percentage of observations within each visually assigned class that were assigned to each predicted class. Performance was evaluated using the same 131 mode-separated tracks, each of which independently met segment-analysis requirements. (a) Point-level performance based on predictions made independently for each dynamic record. (b) Track-level performance after assigning each mode-separated track to the most frequently predicted mode across all records within that track. Overall agreement at the point level was 0.67 and at the track level 0.70.

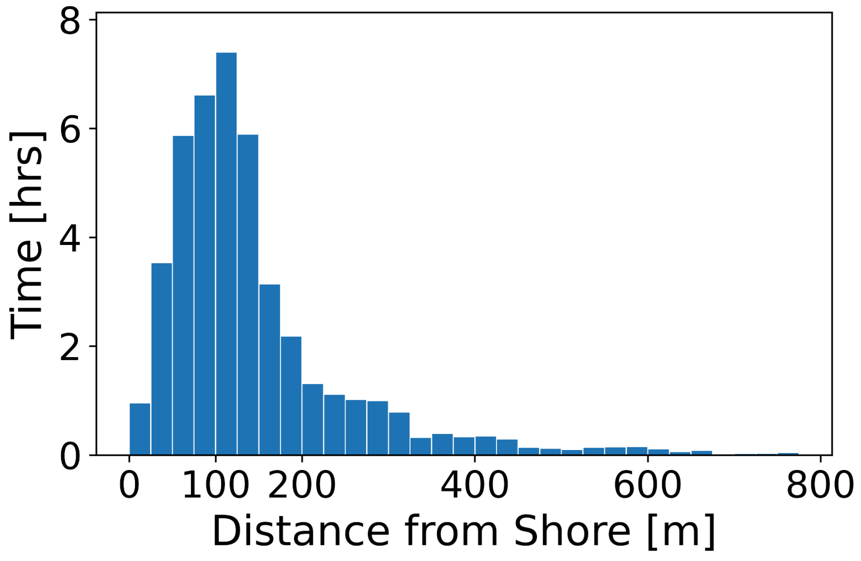


**Figure S12**. Distribution of shark distances from shore for tracks with valid distance-to-shore measurements. Distance-to-shore data were available for 347 of 359 processed tracks. Distances ranged from 3–882 m, with a mean of 145 m and a median of 116 m. Tracking time increased linearly with distance from shore across 25 m bins out to 125 m ($y = 0.06x - 0.87$, $R^2 = 0.93$), then began to decrease.

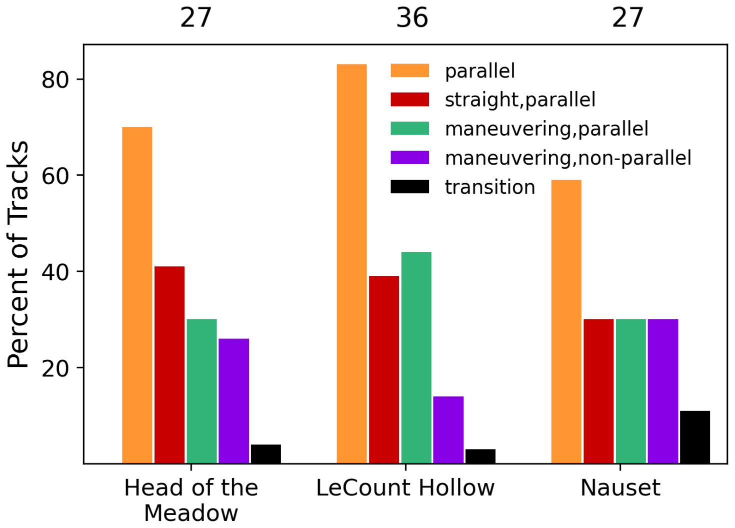


**Figure S13**. Distribution of visually assessed movement modes across the three beach areas for mode-separated tracks derived from initial tracks exceeding 15 min in total duration. Numbers above each beach indicate the number of tracks included. Orange bars show the combined percentage of parallel tracks, including both straight, parallel and maneuvering, parallel modes; the remaining bars show individual movement-mode classifications. LeCount Hollow had a higher ratio of parallel to maneuvering, non-parallel tracks (6.0) than Nauset (2.0) or Head of the Meadow (2.7), although differences among beach areas were not statistically significant.

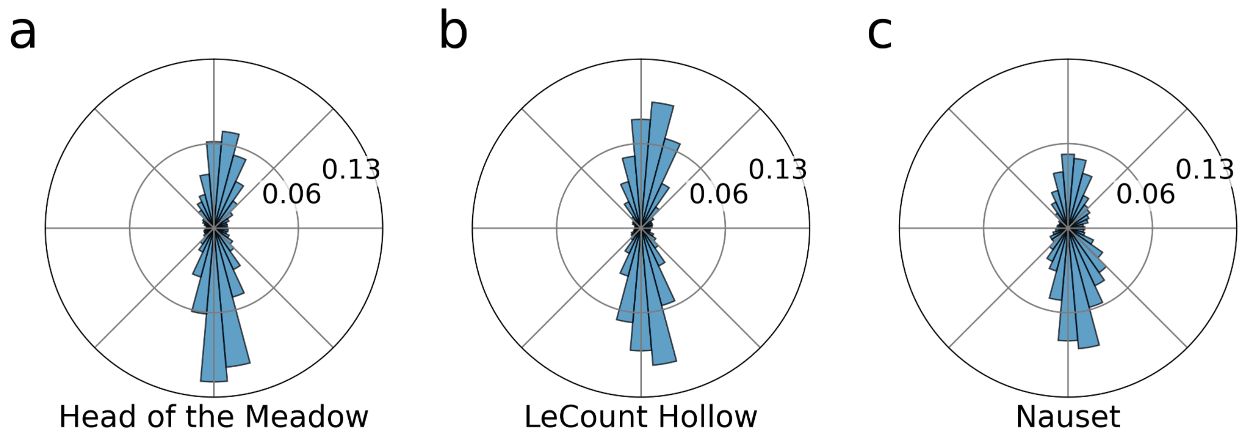


**Figure S14**. Distribution of shark directions of travel relative to the shoreline across the three beach areas. Direction of travel at Nauset (c) was more dispersed from shore-parallel movement than at Head of the Meadow (a) or LeCount Hollow (b). In all plots, bins have a width of 10°, and the radial axis represents the fraction of the total distribution in each bin.

#### Supplementary Tables

**Table S1**. Summary of shark tracks used in segment-based analysis. Initial and mode-separated analytical tracks were independently required to contain at least one continuous segment of at least 5 min with no tracking gaps >30 s. Counts in the minimum-duration columns are based on the total duration of the initial parent track. The number of mode-separated tracks exceeds the number of initial tracks because some initial tracks contained more than one visually assessed movement mode.

| Included Data | Initial Tracks | Mode-Separated Tracks > 5 min | Mode-Separated Tracks > 10 min | Mode-Separated Tracks > 15 min |
|---|---|---|---|---|
| Meeting Segment Requirements | 121 | 131 | 103 | 67 |

**Table S2**. Movement mode assignment based on visual classification. A total of 229 mode-separated tracks derived from initial tracks exceeding 5 min in total duration were categorized into one of four movement modes based on visual assessment. The table reports the count and percentage of the total for each mode. The “Parallel” row is a combined summary category that includes both straight, parallel and maneuvering, parallel classifications and is therefore not an additional mutually exclusive movement mode.

| Movement Mode | Mode Separated Tracks [counts] | Mode Separated Tracks [% of Total] |
|---|---|---|
| Straight, Parallel | 100 | 44 |
| Maneuvering, Parallel | 83 | 36 |
| Maneuvering, Non-Parallel | 30 | 13 |
| Transition | 16 | 7 |
| Parallel | 183 | 80 |
| Total | 229 | 100 |

**Table S3**. Shark ground speeds associated with specific behaviors and visually assessed movement modes. For each behavior or mode, mean ground speed was compared with the mean ground speed across all other observations using Welch’s t-test, with Bonferroni correction applied across comparisons. Mean ground speeds differed significantly for circling, slinky swim, and straight, parallel movement after Bonferroni correction.

| Specific Behavior | Specific Behavior (n, M ± SD [m/s]) | Other Behaviors (n, M ± SD [m/s]) | Welch’s t-test | Bonferroni-corrected p-value |
|---|---|---|---|---|
| Circling | 7, 0.58 ± 0.05 | 206, 1.07 ± 0.26 | t(25.0) = -18.9 | 1.5e-15 |
| Slinky Swim | 5, 0.65 ± 0.06 | 208, 1.06 ± 0.27 | t(8.2) = -12.1 | 1.0e-05 |
| Straight, Parallel | 95, 1.13 ± 0.25 | 118, 0.99 ± 0.27 | t(207.8) = 3.9 | 7.2e-04 |
| Maneuvering, Parallel | 76, 1.03 ± 0.26 | 137, 1.06 ± 0.28 | t(165.4) = -0.68 | 1.00 |
| Maneuvering, Non-Parallel | 20, 0.99 ± 0.27 | 193, 1.06 ± 0.27 | t(23.3) = -1.1 | 1.00 |
| Transition | 10, 1.08 ± 0.22 | 203, 1.05 ± 0.28 | t(10.4) = 0.41 | 1.00 |

**Table S4**. Tailbeat frequencies associated with specific behaviors and visually assessed movement modes. For each behavior or mode, mean tailbeat frequency was compared with the mean tailbeat frequency across all other observations using Welch's t-test, with Bonferroni correction applied across comparisons. Mean tailbeat frequencies differed significantly for circling and slinky swim after Bonferroni correction; no other behaviors or movement modes showed significant differences.

| Specific Behavior | Specific Behavior (n, M ± SD [Hz]) | Other Behaviors (n, M ± SD [Hz]) | Welch's t-test | Bonferroni-corrected p-value |
|---|---|---|---|---|
| Circling | 4, 0.33 ± 0.02 | 75, 0.43 ± 0.09 | t(12.9) = -7.0 | 5.8e-05 |
| Slinky Swim | 4, 0.29 ± 0.04 | 75, 0.43 ± 0.09 | t(4.6) = -6.2 | 0.01 |
| Straight, Parallel | 25, 0.44 ± 0.07 | 54, 0.42 ± 0.10 | t(63.7) = 0.74 | 1.00 |
| Maneuvering, Parallel | 29, 0.44 ± 0.10 | 50, 0.42 ± 0.08 | t(51.5) = 1.3 | 1.00 |
| Maneuvering, Non-Parallel | 11, 0.45 ± 0.10 | 68, 0.42 ± 0.09 | t(12.8) = 0.89 | 1.00 |
| Transition | 6, 0.41 ± 0.05 | 73, 0.43 ± 0.09 | t(7.7) = -0.91 | 1.00 |

**Table S5**. Pairwise chi-square tests with Yates' continuity correction were used to compare movement-mode proportions among beach areas. Comparisons were based on mode-separated tracks derived from initial tracks exceeding 15 min in total duration and classified by visual assessment. For each beach, counts are shown as parallel / maneuvering, non-parallel, where parallel includes both straight, parallel and maneuvering, parallel modes. Transition tracks are not included. While Nauset and Head of the Meadow had lower ratios of parallel to maneuvering, non-parallel tracks as compared to LeCount Hollow, no pairwise comparisons were statistically significant. Site abbreviations are used for compactness: HM = Head of the Meadow and LeCounts = LeCount Hollow.

| Beach 1 (parallel / maneuvering, non-parallel) | Beach 2 (parallel / maneuvering, non-parallel) | $\chi^2$ test p-value |
|---|---|---|
| Nauset (16 / 8) | LeCounts (30 / 5) | 0.157 |
| Nauset (16 / 8) | HM (19 / 7) | 0.853 |
| HM (19 / 7) | LeCounts (30 / 5) | 0.367 |

**Table S6**. Pairwise comparisons of the per-track proportion of time spent traveling within ±30° of the shoreline ("parallel fraction") across beach areas. Reported fractions represent the mean per-track proportion of time parallel to shore. A Kruskal-Wallis test indicated a significant overall difference among beach areas ($H = 12.0$, $p = 2.5 \times 10^{-3}$). Tracks at LeCount Hollow had a significantly greater parallel fraction than tracks at Nauset (Bonferroni-corrected $p = 8.9 \times 10^{-4}$; Cliff's $\delta = -0.413$, medium effect, with the sign reflecting the Nauset–LeCounts comparison order). Other pairwise comparisons were not statistically significant. Site abbreviations are used for compactness: HM = Head of the Meadow and LeCounts = LeCount Hollow.

| Beach 1 (±30° of shoreline) | Beach 2 (±30° of shoreline) | Bonferroni-corrected p-value | Effect Size (Cliff's δ) | Effect Size |
|---|---|---|---|---|
| Nauset (0.57) | LeCounts (0.75) | 8.9e-4 | -0.413 | Medium |
| Nauset (0.57) | HM (0.69) | 0.113 | -0.248 | Small |
| LeCounts (0.75) | HM (0.69) | 0.409 | 0.170 | Small |
